# Label-free Method for Classification of T cell Activation

**DOI:** 10.1101/536813

**Authors:** Alex J. Walsh, Katie Mueller, Isabel Jones, Christine M. Walsh, Nicole Piscopo, Natalie N. Niemi, David J. Pagliarini, Krishanu Saha, Melissa C. Skala

## Abstract

T cells have a range of cytotoxic and immune-modulating functions, depending on activation state and subtype. However, current methods to assess T cell function use exogenous labels that often require cell permeabilization, which is limiting for time-course studies of T cell activation and non-destructive quality control of immunotherapies. Label-free optical imaging is an attractive solution. Here, we use autofluorescence imaging of NAD(P)H and FAD, co-enzymes of metabolism, to quantify optical imaging endpoints in quiescent and activated T cells. Machine learning classification models were developed for label-free, non-destructive determination of T cell activation state. T cells were isolated from the peripheral blood of human donors, and a subset were activated with a tetrameric antibody against CD2/CD3/CD28 surface ligands. NAD(P)H and FAD autofluorescence intensity and lifetime of the T cells were imaged using a multiphoton fluorescence lifetime microscope. Significant differences in autofluorescence imaging end-points were observed between quiescent and activated T cells. Feature selection methods revealed that the contribution of the short NAD(P)H lifetime (*α*_1_) is the most important feature for classification of activation state, across multiple donors and T cell subsets. Logistic regression models achieved 97-99% accuracy for classification of T cell activation from the autofluorescence imaging endpoints. Additionally, autofluorescence imaging revealed NAD(P)H and FAD autofluorescence differences between CD3^+^CD8^+^ and CD3^+^CD4^+^ T cells, and random forest models of the autofluorescence imaging endpoints achieved 97+% accuracy for four-group classification of quiescent and activated CD3^+^CD8^+^ and CD3^+^CD4^+^ T cells. Altogether these results indicate that autofluorescence imaging of NAD(P)H and FAD is a powerful method for label-free, non-destructive determination of T cell activation and subtype, which could have important applications for the treatment of cancer, autoimmune, infectious, and other diseases.

## 1 Introduction

T cells are an important component of the adaptive immune response and have diverse cytotoxic and immune-modulating, or “helper” activities, upon activation. The two main T cell subtypes are CD3^+^CD8^+^ T cells that engage in cell-mediated cytotoxicity and release toxic cytokines, including interferon gamma (IFN-*γ*) and tumor necrosis factor alpha (TNF-*α*), and CD3^+^CD4^+^ T cells that can be further divided into additional subtypes with differing pro- and anti-inflammatory functions due to chemokine and cytokine production[1, 2]. T cells are a promising target for immunotherapies because of these diverse functions. Immunotherapies that directly increase T cell cytotoxic activity, such as immune checkpoint blockade therapies and adoptive cell transfer therapies, are currently used clinically for cancer treatment and are in development for additional diseases including HIV[3, 4]. Immunotherapies that enhance regulatory T cell (T_*REG*_) behaviors are in 42 development to treat transplant rejection and autoimmune diseases, including diabetes and Crohn’s disease [5–7]. Due to the variable behaviors of T cell subsets, full evaluation of immunotherapy efficacy requires profiling of T cell subtypes and activation states to assess the impact of different T cell compartments on the patient, select for appropriate therapeutic cell populations, and evaluate the degree of response upon stimulation.

New tools that are non-destructive and label-free are needed to fully characterize T cells for assessment of immunotherapies. Currently, T cell subtype and function is determined from expression of surface proteins (e.g. CD3, CD4, CD8, CD45RA, etc.) and cytokine production (e.g. IFN-*γ*, TGF-*β*, IL-2, IL-4, IL-17, etc.) by antibody-based methods such as flow cytometry, immunohistochemistry, or immunofluorescence, or by transgenic fluorophore expression. However, all of these methods require exogenous contrast agents, and flow cytometry and immunohistochemistry require tissue dissociation and fixation, respectively. A non-destructive 5 and label-free method of determining T cell activity would enable direct observation of T cell behavior and immunotherapy effects *in vivo* in preclinical models of cancer. Additionally, such a tool could be amenable 55 for single-cell quality control of adoptive T cell therapies, where T cells, expanded *in vitro*, are injected into the patient. Autofluorescence imaging is an attractive method to probe immune cell behaviors because it is non-destructive, relies on endogenous contrast, and provides high spatial and temporal resolution.

Fluorescence imaging of the endogenous metabolic co-enzymes NAD(P)H and FAD provides quantitative endpoints of cellular metabolism [8–10]. (NADH and NADPH fluorescence are indistinguishable; therefore, NAD(P)H is used to represent the combined fluorescence signal[11].) The optical redox ratio is the fluorescence intensity of NAD(P)H divided by the sum of the fluorescence intensities of NAD(P)H and FAD, and an optical measurement of the redox state of the cell [8, 12]. The fluorescence lifetime, the time the fluorophore is in the excited state before returning to ground state, provides information on the protein binding of NAD(P)H and FAD [9, 13]. NAD(P)H and FAD can both exist in two conformations: a quenched and unquenched form, with a short and long lifetime, respectively. NAD(P)H has a short lifetime in the free state and a long lifetime in its protein-bound state [9]. Conversely, FAD has a short lifetime when bound to an enzyme and a long lifetime when free [13]. Fluorescence lifetime imaging (FLIM) allows quantification of the short (*τ*_1_) and long (*τ*_2_) lifetime values, the fraction of free and protein-bound co-enzyme (*α*_1_ and *α*_2_, respectively, for NAD(P)H, and *α*_2_ and *α*_1_, respectively, for FAD), and the mean lifetime (the weighted average of the short and long lifetimes, *τ*_*m*_ = *α*_1_ ∗ *τ*_1_ + *α*_2_ ∗ *τ*_2_). The fluorescence intensity and lifetime of NAD(P)H and FAD are sensitive to metabolic differences between neoplasias and malignant tis-2 sues, anti-cancer drug effects in cancer cells, and differentiating stem cells [14–19]. Autofluorescence imaging has been used previously to identify macrophages *in vivo* and detect metabolic changes due to macrophage polarization [20–22]. Altogether, fluorescence lifetime imaging of NAD(P)H and FAD provide quantitative and functional endpoints of cellular metabolism.

T cells undergo metabolic reprogramming when activated by an antigen. Upon activation, T cells have increased metabolic demands to support cell growth, proliferation, and differentiation [23]. CD28 stimulation induces glucose uptake and glycolysis in T cells through upregulation of GLUT1, phosphatidylinositol 3’-9 kinase (PI3K), and Akt. This metabolic state of increased aerobic glycolysis is required for T cells to maintain effector function [23–25]. Therefore, this study tests the hypothesis that fluorescence lifetime imaging of NAD(P)H and FAD provides a label-free, non-destructive method with quantitative endpoints to identify activated T cells. To test this hypothesis, we isolated T cells from the blood of healthy donors, activated the cells in an antigen-independent manner with a tetrameric antibody (anti-CD2/CD3/CD28) and imaged the NAD(P)H and FAD fluorescence intensity and lifetime of quiescent and activated T cells. This is the first study to (1) demonstrate autofluorescence lifetime differences between quiescent and activated T cells and (2) accurately classify T cell activation state from machine learning models using quantitative endpoints from autofluoresence lifetime images.

## 2 Results

### 2.1 Autofluorescence imaging reveals metabolic differences with activation in T cells

T cell isolations for CD3^+^ (pan-T cell marker) and CD3^+^CD8^+^ cells were used to study all T cells, as might be utilized in adoptive cell transfer therapies, and the cytotoxic CD3^+^CD8^+^ sub-population, respectively. NAD(P)H and FAD autofluorescence imaging reveals metabolic differences in quiescent and activated T cells (Fig. 1, S1). The high resolution multiphoton imaging allows visualization of bulk CD3^+^ and isolated CD3^+^CD8^+^ T cells (Fig. 1A). In the autofluorescence images, the nucleus remains dark as NAD(P)H is primarily located in the cytoplasm and mitochondria, and FAD is primarily in the mitochondria. Immunoflu-7 orescence labeling of CD4, CD8, and CD69 surface proteins verified cell type and activation (Fig. S2). There were significant differences in cell size, optical redox ratio, NAD(P)H *τ*_*m*_, NAD(P)H *α*_1_, and FAD *α*_1_ between quiescent and activated T cells (p<0.001, Fig. 1B-F). Significant changes (p<0.001) in FAD *τ*_*m*_ between quiescent and activated T cells were found only for T cells within the bulk CD3^+^ T cell population (Fig. 1E). Additionally, significant changes (p<0.001) in the short and long lifetimes were observed between quiescent and activated CD3^+^ and CD3^+^CD8^+^ T cells (Fig. S1). These differences in autofluorescence endpoints were consistent across the 6 donors (Fig. 1, S1), at 24 and 48 hr of exposure to the activating antibodies (Fig. S3), and between experiments from two different blood draws (183 days apart) from the same donor (Fig. S4). A slight increase in FAD *τ*_1_ was found in both quiescent and activated CD3^+^ T cells, suggesting 106 a slight change in the microenvironment of bound FAD between CD3^+^ T cells of the same donor from two blood draws; however, no other autofluorescence endpoints were signficantly different between the two blood draws.

**Figure 1:**
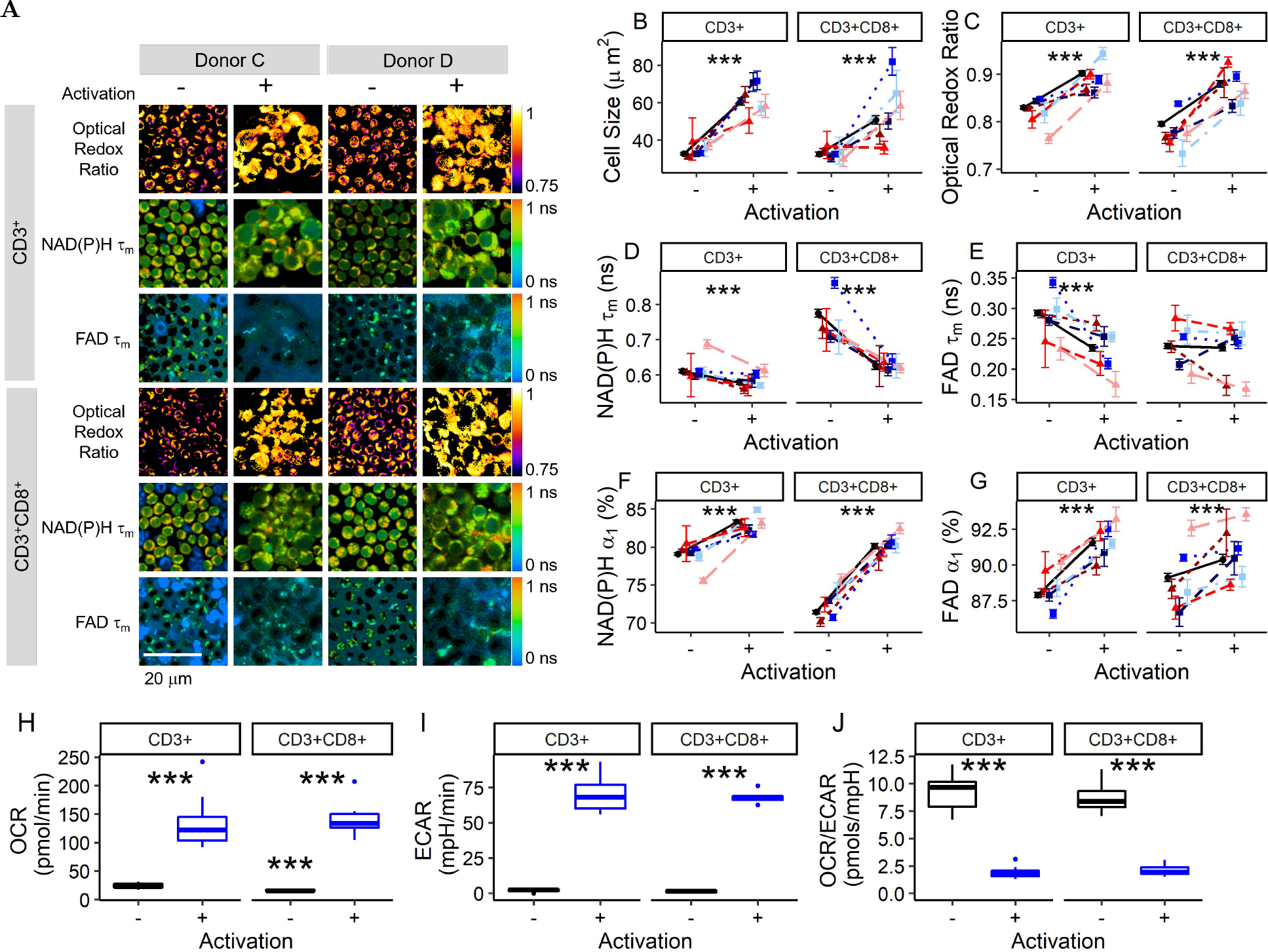
NAD(P)H and FAD autofluorescence imaging reveals metabolic differences between quiescent and activated T cells. Representative optical redox ratio, NAD(P)H *τ*_*m*_, and FAD *τ*_*m*_ images of quiescent (columns 1, 3) and activated (columns 2, 4) CD3^+^ (rows 1-3) and CD3^+^CD8^+^ (row 4-6) T cells from two different donors. Scale bar is 20 *µ*m. Cell size (B), optical redox ratio (C), NAD(P)H *τ*_*m*_ (D), FAD *τ*_*m*_ (E), NAD(P)H *α*_1_ (F), and FAD *α*_1_ (G) of quiescent and activated CD3^+^ and CD3^+^CD8^+^ T cells. Black circles represent mean of all data (6 donors), triangles (donors A [dark red], B [medium red], and F [light red]) represent data from female donors, squares (donors C [dark blue], D [medium blue], and E [light blue]) represent data from male donors. Each color shade represents data from an individual donor. Data are mean +/− 99% CI. *** p<0.001. n = 54-1058 cells per donor per group. (H-J) Cellular respiration increases in activated T cells. The oxygen consumption rate (OCR; panel H) and extracellular acidification rate (ECAR, panel I) are increased in activated bulk CD3^+^ and isolated CD3^+^CD8^+^ T cells. The ratio of OCR to ECAR (J) is significantly decreased in activated bulk CD3^+^ and isolated CD3^+^CD8^+^ T cells as compared with that of quiescent T cells. *** p<0.001, Student’s t-test, n = 6 wells/group CD3^+^CD8^+^ isolation, n = 12 wells/group CD3^+^ isolation.

Seahorse OCR and ECAR measurements confirm increased metabolic rates of the activated T cells (p<0.001, Fig. 1H-J). In a metabolic inhibitor experiment (Fig. S5), the redox ratio of activated T cells decreased (p<0.001) with a glycolysis inhibitor (2-deoxy-d-glucose), and the redox ratio of quiescent T cells increased (p<0.001) with oxidative phosphorylation inhibitors (antimycin A and rotenone). Additionally, the glutaminolysis inhibitor BPTES significantly decreased (p<0.001) the optical redox ratio, NAD(P)H *τ*_*m*_, and FAD *τ*_*m*_ of both quiescent and activated T cells, suggesting a significant contribution of glutaminolysis to the metabolism of quiescent and activated T cells (Fig. S5).

### 2.2 Machine learning models of autofluorescence imaging endpoints allow classification of quiescent and activated T cells with high accuracy

Uniform Manifold Approximate and Projection (UMAP) [26], a dimension reduction technique similar to tSNE, was used to visualize how cells cluster from autofluorescence measurements. Neighbors were defined through a cosine distance function computed across the autofluorescence endpoints (optical redox ratio, NAD(P)H *τ*_*m*_, NAD(P)H *τ*_1_, NAD(P)H *τ*_2_, NAD(P)H *α*_1_, FAD *τ*_*m*_, FAD *τ*_1_, FAD *τ*_2_, and FAD *α*_1_) and cell size for each cell. UMAP was chosen over other techniques, notably PCA or tSNE, for its speed, ability to include non-metric distance functions, and performance on preserving the global structure of the data. UMAP representations of the autofluorescence imaging data reveals separation of quiescent and activated T cells (Fig. 2A-B). The gain ratio of autofluorescence endpoints indicates that NAD(P)H *α*_1_, cell size, and optical redox ratio are the most important features for classification of activation state of CD3^+^ T cells (Fig. 2C), and NAD(P)H *α*_1_, optical redox ratio, and NAD(P)H *τ*_*m*_ are the most important features for classification of activation state of CD3^+^CD8^+^ T cells (Fig. 2C). The order of feature importance was consistent across multiple feature selection methods including information gain, *χ*^2^, and random forest (Fig. S6). Correlation analysis revealed that NAD(P)H *α*_1_, cell size, and the optical redox ratio are not significantly correlated (Fig. S7), suggesting these features are independent and provide complementary information for classification. NAD(P)H *α*_1_ and *τ*_*m*_ are significantly correlated (Fig. S7), as expected, given that *τ*_*m*_ is computed from *α*_1_. Similar feature weight and order of importance were observed from analysis without NAD(P)H *τ*_*m*_ and FAD *τ*_*m*_ (Fig. S8), indicating that the multivariate models were not significantly affected by the correlations between the mean lifetimes and the lifetime components.

**Figure 2:**
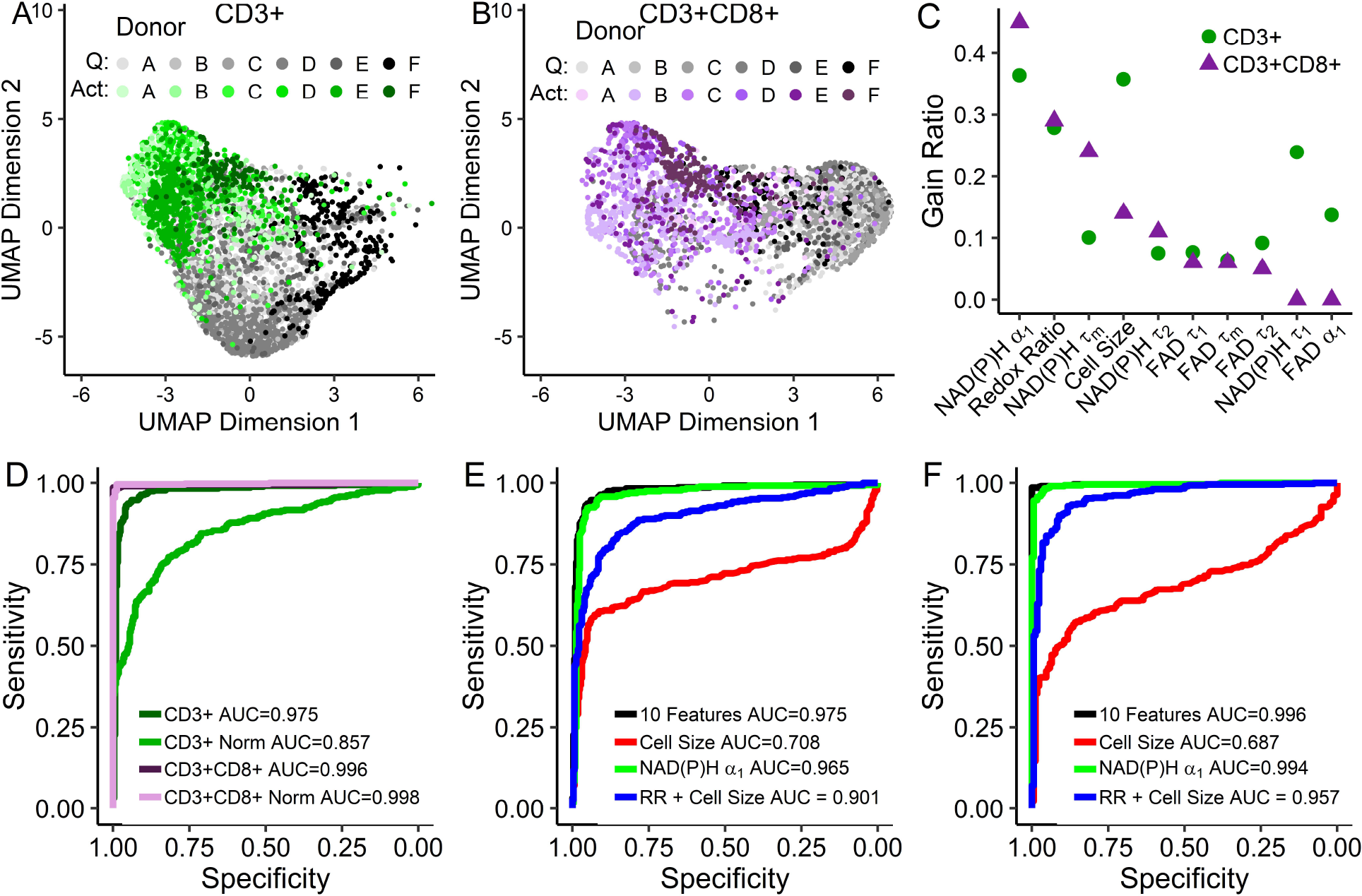
Autofluorescence imaging endpoints allow classification of quiescent and activated T cells. (A-B) UMAP data reduction technique allows visual representation of the separation between quiescent (”Q”) and activated (”Act”) bulk CD3^+^ (A) and isolated CD3^+^CD8^+^ (B) T cells. Each color shade corresponds to a different donor, grays correspond to quiescent cells and green or purple to activated CD3^+^ or CD3^+^CD8^+^ T cells, respectively. (C) Feature weights for classification of quiescent versus activated T cells by the gain ratio method. (D) ROC curves for logistic regression models for classification of activation state within bulk CD3^+^ T cells, bulk CD3^+^ T cells normalized within each donor (CD3^+^ Norm), isolated CD3^+^CD8^+^ T cells, and isolated CD3^+^CD8^+^ T cells normalized within each donor (CD3^+^CD8^+^ Norm). (E-F) ROC curves for logistic regression classification models computed using different features for the classification of (E) quiescent or activated bulk CD3^+^ or (F) isolated CD3^+^CD8^+^ T cells. Models were trained on cells that lacked same cell validation data from donors A, B, C, and D but were known to be quiescent or activated by culture conditions (n = 4131 CD3^+^ cells, n = 2655 CD3^+^CD8^+^ cells), and cells from donors B, E, and F with CD69 validation of activation state were used to test the models (n = 696 CD3^+^ cells, n = 595 CD3^+^CD8^+^ cells).

Classification models were developed to predict T cell activation state from NAD(P)H and FAD autofluorescence imaging endpoints (Fig. 2D-F). To protect against over-fitting, models were trained on data from 4 donors with activation state assigned from culture conditions and tested on data with same-cell CD69 expression immunofluorescence validation from 3 donors (completely independent and non-overlapping ob-40 servations). Receiver operator characteristic (ROC) curves reveal high classification accuracy for predicting activation in bulk CD3^+^ (AUC = 0.975) and isolated CD3^+^CD8^+^ (AUC = 0.996) T cells, when the models use all autofluorescence endpoints (optical redox ratio, cell size, NAD(P)H *τ*_*m*_, NAD(P)H *τ*_1_, NAD(P)H *τ*_2_, NAD(P)H *α*_1_, FAD *τ*_*m*_, FAD *τ*_1_, FAD *τ*_2_, and FAD *α*_1_). When the NAD(P)H and FAD autofluorescence imaging endpoints of the T cells are normalized within a donor to the mean value of the quiescent CD3^+^ population, the ROC AUC decreases to 0.857 for CD3^+^ T cells (Fig. 2D) and increases slightly to 0.998 for isolated CD3^+^CD8^+^ T cells. While all 10 NAD(P)H and FAD autofluorescence features achieved the highest classification accuracy (AUC = 0.975) for activation of CD3^+^ T cells, a model using only NAD(P)H *α*_1_ achieved a slightly lower accuracy of 0.965 (Fig. 2E). Models that include cell size or cell size and the optical redox ratio, endpoints that can be obtained from fluorescence intensity images, were less effective at accurately predicting activation of bulk CD3^+^ T cells with ROC AUCs of 0.708 and 0.901, respectively (Fig. 2E). Similar results were obtained for the isolated CD3^+^CD8^+^ T cells, with the highest ROC AUC values achieved for logistic regression classification models using all 10 autofluorescence imaging endpoints and NAD(P)H *α*_1_ alone, AUC = 0.996 and 0.994, respectively (Fig. 2F). Similar classification accuracy was achieved with random forest and support vector machine models using all 10 autofluorescence imaging endpoints (Fig. S9).

### 2.3 Autofluorescence imaging reveals T cell heterogeneity within and across donors

T cell heterogeneity was assessed within and across donors (Fig. 3). Heatmap representation (Fig. 3A) of the z-score of autofluorescence imaging endpoint values at the donor level (each row is the mean data of a single donor, cell type, and activation) reveals that the T cells cluster by activation state (i.e. quiescent and activated cluster separately) and isolation (bulk CD3^+^ or isolated CD3^+^CD8^+^). Corresponding coefficient of variation heatmaps highlight the high intra-donor variability of the size of activated T cells and low intra-donor heterogeneity of the autofluorescence endpoints (Fig. S10).

**Figure 3:**
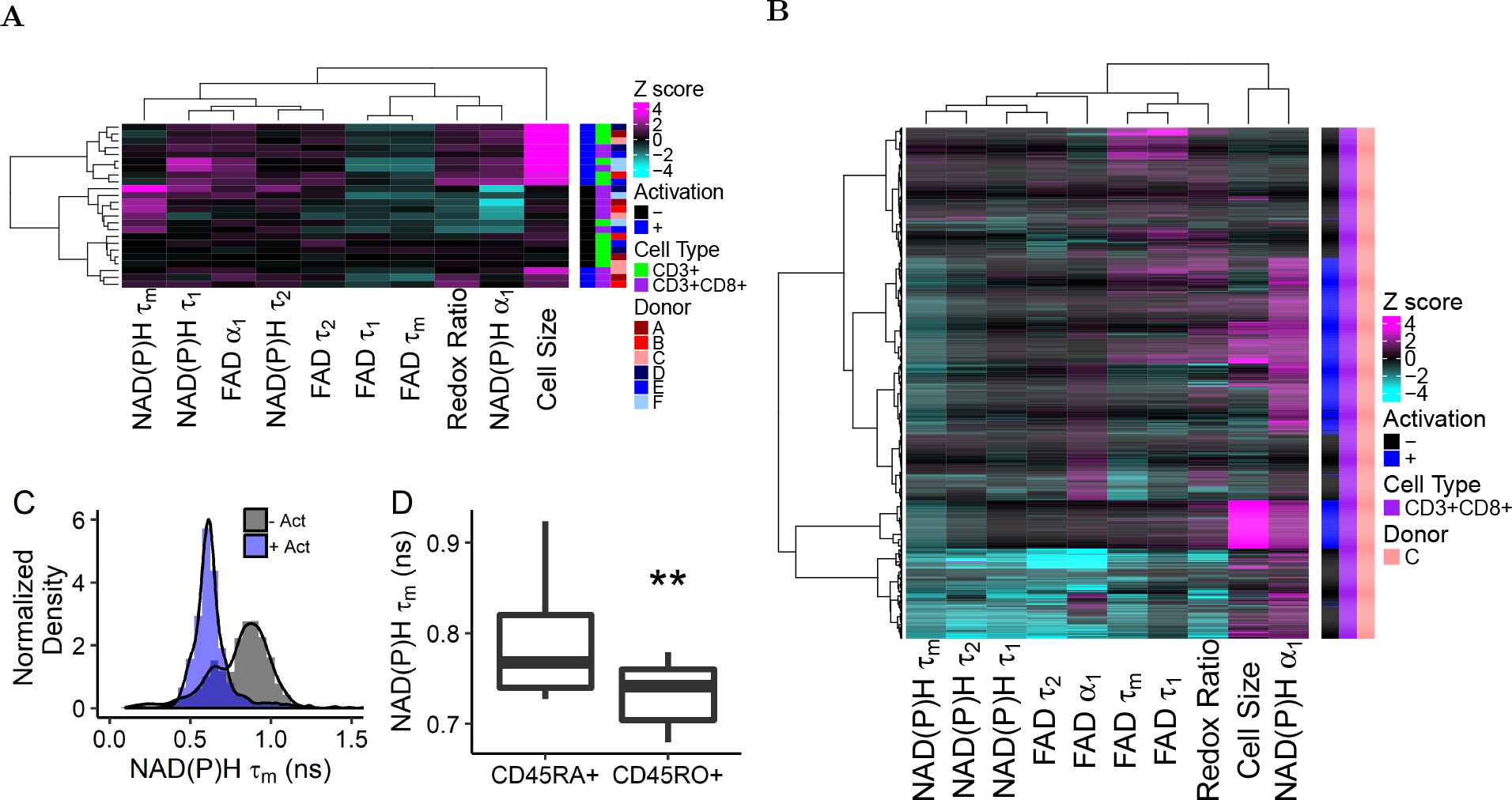
Autofluorescence imaging reveals inter- and intra-donor T cell heterogeneity. (A) Heatmap of z-scores of NAD(P)H and FAD autofluorescence imaging endpoints where each row is the mean data representing a single donor, subtype (CD3^+^ or CD3^+^CD8^+^), and activation. Data clusters by activation state and isolation (bulk CD3^+^ or isolated CD3^+^CD8^+^). (B) Heatmap of z-scores of NAD(P)H and FAD autofluorescence imaging endpoints of CD3^+^CD8^+^ T cells from a single donor, each row is a single cell (n=635 cells). Distinct clusters are identified within the quiescent and activated CD3^+^CD8^+^ T cells. (C) Histogram analysis of NAD(P)H *τ*_*m*_ reveals two populations in quiescent CD3^+^CD8^+^ T cells across all donors (n=2126 quiescent cells, 1352 activated cells). (D) NAD(P)H *τ*_*m*_ is decreased in CD45RO+ CD3^+^CD8^+^ T cells compared to NAD(P)H *τ*_*m*_ of CD45RA+ CD3^+^CD8^+^ T cells (CD45RA+ n=27 cells, CD45RO+ n=11 cells from 1 donor, ** p<0.01, - Act = quiescent cells, + Act = cells exposed to anti-CD3/CD2/CD28 for 48hr.)

A representative z score heatmap where each row is a single cell from one donor reveals distinct clusters of T cells by autofluorescence imaging endpoints within the quiescent and activated CD3^+^CD8^+^ T cell populations (Fig. 3B). Multiple quiescent and activated T cell populations were observed across all six donors and arises from varied distributions of autofluorescence imaging endpoints within the T cell populations (Fig. 3C, S11-12). For example, histograms of the NAD(P)H *τ*_*m*_ values of quiescent and activated CD3^+^CD8^+^ T cells reveals a bimodal population within the quiescent CD3^+^CD8^+^ T cells, with one peak of the quiescent cells consistent with the peak of the activated cells (Fig. 3C).

We hypothesized that memory and naïve T cells within the quiescent population contributed to the observed heterogeneity within the quiescent CD3^+^CD8^+^ T cell population (Fig. 3B-C, S11-13) (i.e. the multiple clusters of quiescent CD3^+^CD8^+^ cells within the heatmaps and bimodal distribution of the NAD(P)H *τ*_*m*_ of quiescent CD3^+^CD8^+^ T cells). To test this, we co-stained quiescent CD3^+^CD8^+^ T cells with antibodies against CD45RA, a marker of naïve T cells, and CD45RO, a marker of memory T cells. NAD(P)H *τ*_*m*_ was significantly decreased in CD45RO+ cells as compared with NAD(P)H *τ*_*m*_ of CD45RA+ cells (Fig. 3D). Additionally, the optical redox ratio and NAD(P)H *α*_1_ were increased (p<0.01) in CD45RO+ CD3^+^CD8^+^ T cells as compared to CD45RA+ cells (Fig. S14).

### 2.4 Culture with CD3^+^CD4^+^ T cells affects the autofluorescence of CD3^+^CD8^+^ T cells

NAD(P)H and FAD autofluorescence imaging endpoints reveal metabolic differences between CD3^+^CD8^+^ T cells cultured as an isolated population and CD3^+^CD8^+^ T cells cultured with CD3^+^CD4^+^ T cells (bulk CD3^+^ isolation). A UMAP (data dimension reduction) representation of NAD(P)H and FAD autofluorescence imaging endpoints reveals that CD3^+^CD8^+^ T cells cultured from the CD3^+^CD8^+^ specific T cell isolations cluster separately from CD3^+^CD8^+^ T cells within bulk CD3^+^ T cell populations (Fig. 4A). The optical redox ratio and NAD(P)H *α*_1_ are decreased in both quiescent and activated CD3^+^CD8^+^ T cells of the isolated CD3^+^CD8^+^ population as compared to the corresponding values of quiescent and activated CD3^+^CD8^+^ T cells, respectively, within the bulk CD3^+^ population (Fig. 4B-C). Additional differences in NAD(P)H and FAD autofluorescence lifetime endpoints were observed between CD3^+^CD8^+^ T cells within the bulk CD3^+^ population and the isolated CD3^+^CD8^+^ population (Fig. S15).

**Figure 4:**
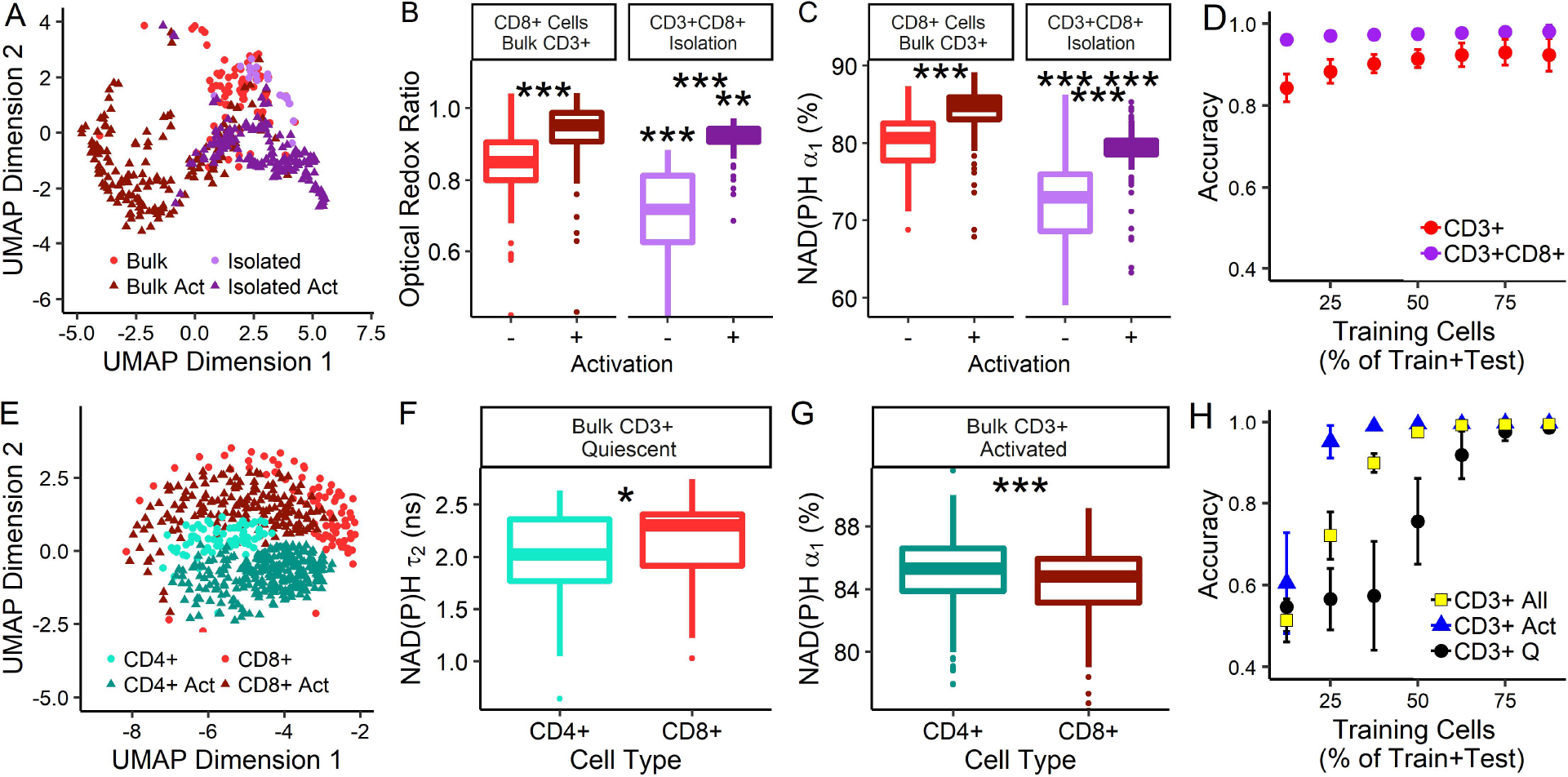
T cell population composition affects T cell autofluorescence. (A) UMAP of NAD(P)H and FAD autofluorescence endpoints of quiescent and activated (“Act”) CD3^+^CD8^+^ T cells identified within bulk CD3^+^ and specific CD3^+^CD8^+^ isolations. (B) Optical redox ratio and (C) NAD(P)H *α*_1_ of CD3^+^CD8^+^ T cells cultured as an isolated population (CD3^+^CD8^+^ specific isolation, n=39 quiescent cells, n=174 activated cells) and with CD3^+^CD4^+^ T cells (bulk CD3^+^ isolation, n=83 quiescent cells, n=170 activated cells). Stars between quiescent and activated boxplots compare quiescent and activated CD3^+^CD8^+^ T cells within an isolation (CD3^+^ or CD3^+^CD8^+^), stars above the quiescent box plot represent signficance between quiescent CD3^+^CD8^+^ T cells from the bulk CD3^+^ and CD3^+^CD8^+^ specific isolations, stars above the activated box plot represent signficance between activated CD3^+^CD8^+^ T cells from the bulk CD3^+^ and CD3^+^CD8^+^ specific isolations, ** p<0.01, *** p<0.001. (D) Accuracy of random forest classification of quiescent versus activated CD3^+^CD8^+^ T cells from CD3^+^CD8^+^ specific isolations (n=213 cells) and bulk CD3^+^ isolations (n=253 cells). (E) UMAP of NAD(P)H and FAD autofluorescence imaging endpoints of quiescent and activated CD3^+^CD4^+^ and CD3^+^CD8^+^ cells identified within bulk CD3^+^ populations. (F) NAD(P)H *τ*_2_ of quiescent CD3^+^CD4^+^ and CD3^+^CD8^+^ cells (bulk CD3^+^ isolation, n=66 quiescent CD3^+^CD4^+^ T cells, n=83 quiescent CD3^+^CD8^+^ T cells, * p<0.05, *** p<0.001). (G) NAD(P)H *α*_1_ of activated CD3^+^CD4^+^ and CD3^+^CD8^+^ cells (bulk CD3^+^ isolation, n=264 activated CD3^+^CD4^+^ T cells, n=170 activated CD3^+^CD8^+^ T cells). (H) Accuracy of random forest classification of CD3^+^CD4^+^ and CD3^+^CD8^+^ T cells from quiescent (2 group classification, “CD3^+^ Q”), activated (2 group classification, “CD3^+^ Act”), or both quiescent and activated T cells (4 group classification, “CD3^+^ All”) within bulk CD3^+^ isolations, total observations include 66 quiescent CD3^+^CD4^+^ T cells, 83 quiescent CD3^+^CD8^+^ T cells, 264 activated CD3^+^CD4^+^ T cells, and 170 activated CD3^+^CD8^+^ T cells.

Despite these differences between CD3^+^CD8^+^ T cells of CD3^+^CD8^+^ specific isolations and bulk CD3^+^ isolations, significant changes in NAD(P)H and FAD autofluorescence endpoints due to activation are main-93 tained, and classification models predict activation status of CD3^+^CD8^+^ cells with high accuracy regardless of isolation (Fig. 4D). Random forest feature selection revealed that NAD(P)H *α*_1_ is the most important feature for classification of quiescent from activated CD3^+^ or CD3^+^CD8^+^ T cells (Fig. S16A).

### 2.5 Machine learning models of autofluorescence endpoints classify CD3^+^CD4^+^ from CD3^+^CD8^+^ T cells within bulk CD3^+^ populations

Heterogeneity in NAD(P)H and FAD autofluorescence endpoints between CD3^+^CD4^+^ and CD3^+^CD8^+^ T cells was observed within the T cells from the bulk CD3^+^ isolation. A UMAP representation of the NAD(P)H and FAD autofluorescence data allows visualization of the clustering and separation of quiescent and activated CD3^+^CD4^+^ and CD3^+^CD8^+^ T cells within the bulk CD3^+^ isolation (Fig. 4E). These differences between CD3^+^CD4^+^ and CD3^+^CD8^+^ T cells are due to significant differences in NAD(P)H and FAD endpoints, including NAD(P)H *τ*_2_, which is increased (p<0.05) in quiescent CD3^+^CD8^+^ T cells compared to quiescent CD3^+^CD4^+^ T cells, and NAD(P)H *α*_1_, which is decreased in activated CD3^+^CD8^+^ T cells compared to activated CD3^+^CD4^+^ T cells (p<0.05, Fig. 4F-G, S17). Random forest models to classify T cell subtype (CD3^+^CD4^+^ or CD3^+^CD8^+^) within the bulk CD3^+^ T cell isolation have average predictions of 97.5% and 99.7% for separate predictions on subsets of quiescent or activated T cells, respectively, and 99.4% for all four groups, when trained on 75% of the T cell observations and tested on the remaining 25% (Fig. 4H). Classification accuracy scales with number of cells in train versus test groups (Fig. 4H). Random forest feature analysis revealed that NAD(P)H *τ*_2_ is the highest weighted feature for the classification of activated CD3^+^CD4^+^ from activated CD3^+^CD8^+^ T cells, and FAD *τ*_1_ is the highest weighted feature for quiescent CD3^+^CD4^+^ from quiescent CD3^+^CD8^+^ T cells (Fig. S16B).

### 2.6 Autofluorescence imaging allows classification of activated T cells in cultures of combined quiescent and activated T cells

NAD(P)H and FAD autofluorescence imaging allows label-free imaging and classification of T cell activation in T cell cultures with combined quiescent and activated cells. A representative NAD(P)H *α*_1_ image with CD69 immunofluorescence overlaid in pink, demonstrates the difference in NAD(P)H *α*_1_ between quiescent (CD69^−^) and activated (CD69^+^) T cells (Fig. 5A). UMAP visualization of the autofluorescence imaging data reveals separation of quiescent and activated CD3^+^ T cells within this population of combined quiescent and activated cells (Fig. 5B). When cultured in isolated populations, quiescent and activated T cells have significantly different NAD(P)H and FAD imaging endpoints, including the optical redox ratio and NAD(P)H *α*_1_, than their respective counterpart from a combined (quiescent with activated T cells) population (Fig. 5C-D, Fig. S18). Random forest feature selection for classification of activation status of T cells within a combined, quiescent and activated, T cell population reveals that NAD(P)H *α*_1_ is the most important feature for classification, followed by NAD(P)H *τ*_*m*_ (Fig. S19). Logistic regression models to predict activation status of T cells in a combined, quiescent and activated, CD3^+^ T cell culture achieves ROC AUCs of 0.95 when all 10 NAD(P)H and FAD imaging endpoints are included, 0.95 and 0.68 when only predicting from NAD(P)H *α*_1_ or cell size, respectively, and 0.67 for redox ratio and cell size (Fig. 5E).

**Figure 5:**
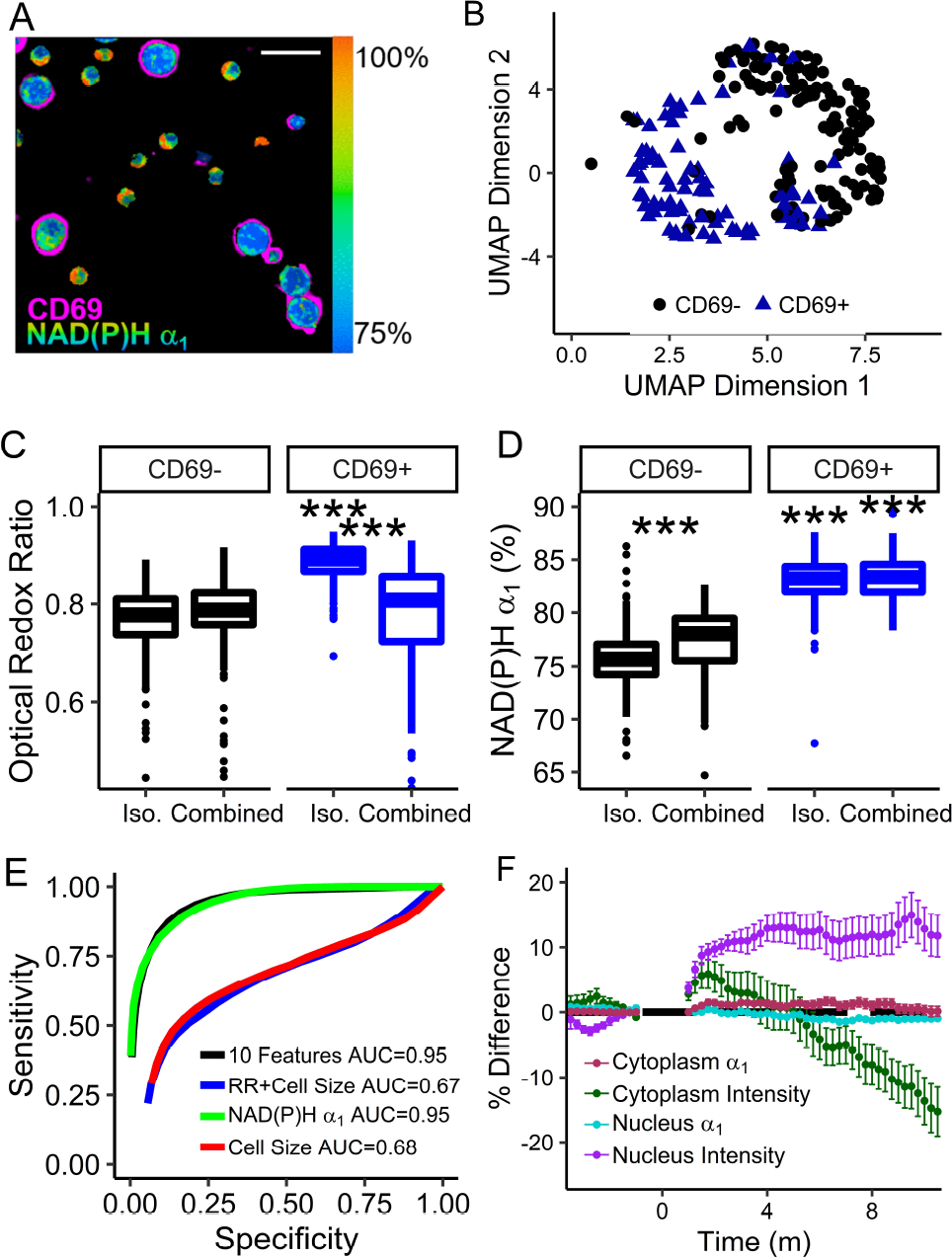
Autofluorescence imaging allows classification of quiescent and activated T cells within combined quiescent and activated T cell populations. (A) Representative NAD(P)H *α*_1_ image of combined quiescent (CD69^−^) and activated (CD69+) T cells with CD69 immunofluorescence overlaid in pink. Scale bar is 30 μm. (B) UMAP representation of NAD(P)H and FAD imaging endpoints of CD69^−^ and CD69+ CD3^+^ T cells from a combined population of quiescent and activated T cells. (C) Optical redox ratio and (D) NAD(P)H *α*_1_ of isolated (“Iso.”) and combined quiescent (CD69^−^) and activated (CD69+) CD3^+^ T cells. *** p < 0.001, n=289-438 cells per group, single donor. (E) ROC curves of logistic regression classification of quiescent and activated CD3^+^ T cells from a combined population of CD69^−^ and CD69+ T cells. (F) Percent difference of NAD(P)H *α*_1_ and fluorescence intensity in CD3^+^ T cell nuclei and cytoplasms over time. Anti-CD2/CD3/CD28 added at t=0 m. mean +/− SD of 34 cells.

### 2.7 Autofluorescence imaging resolves temporal changes in T cells with activation

Metabolic changes occur rapidly within T cells upon activation [27]; therefore, we hypothesized that time-course imaging of T cells would resolve changes in T cell autofluorescence. NAD(P)H fluorescence lifetime images were acquired from CD3^+^ quiescent T cells immediately after exposure to the activating tetrametic antibody (anti-CD2/CD3/CD28). The NAD(P)H intensity of the nucleus increased by 10% relative to the pre-activator values, within a few minutes of addition of the activator, and remained consistently higher than the average pre-activation NAD(P)H intensity throughout the time-course (Fig. 5F). NAD(P)H intensity within the nucleus may indicate increased transcription [28]. The NAD(P)H intensity in the cytoplasm initially increased (t<1 m) and then decreased, relative to the pre-activation NAD(P)H intensity of the cytoplasm. NAD(P)H *α*_1_ increased significantly in the cytoplasm by 2% at t = 6 minutes post addition of the activator and remained significantly increased until t=8.75 m. These autofluorescence changes observed early, within minutes of activation, indicate that autofluorescence lifetime imaging is sensitive to robust transcription and metabolic changes that occur with activation in T cells [27].

## 3 Discussion

T cells are an important component of the adaptive immune response with direct cytotoxic and immune-modulating behaviors. Novel immunotherapies that directly modify T cell behavior show promise for treating a variety of conditions including cancer and autoimmune disease. Due to their varied activities, characterization of T cell function is imperative for assessment of immunotherapy efficacy for pre-clinical evaluation and quality control of clinical immunotherapies. In this study, we develop autofluorescence lifetime imaging-based methods for determination of T cell activation at the single cell level. Autofluorescence lifetime imaging is non-destructive, label-free, and has high spatial and temporal resolution that is amenable with live cell assessment, longitudinal studies, and *in vivo* imaging. Autofluorescence imaging offers advantages over antibody-labeling methods that are traditionally used to assess T cell function with high specificity which are less amenable to non-invasive time-course studies within intact samples.

Upon activation, T cell metabolism switches from tricarboxylic acid oxidation of glucose and *β*-oxidation of fatty acids to glycolysis and glutaminolysis [23–25, 29, 30]. T cells with high glycolytic activity *in vitro* show poor persistence, low recall responses and low proliferation rates that lead to poor effector activity *in vivo*, whereas T cells with high fatty acid oxidation show increased persistence, recall responses and proliferation leading to better effector activity within the tumor [31]. Changes in NAD(P)H and FAD autofluorescence imaging endpoints, including the increased optical redox ratio observed in activated T cells relative to the optical redox ratio of quiescent T cells, reflect a shift towards glycolysis in activated T cells (Fig. 1, S1, S18-19). Significant changes in the lifetimes of protein-bound NAD(P)H (*τ*_2_) and protein-bound FAD (*τ*_1_; Fig. S1) indicate differences in the protein binding partners of NAD(P)H and FAD [32]. A significant increase in the fraction of free NAD(P)H (*α*_1_; Fig. 1F) in activated T cells as compared to that of quiescent T cells, suggests a relative increase in free NAD(P)H and a decrease in protein-bound NADH, consistent with a shift from TCA metabolism to glycolysis [33], which was verified by the Seahorse assay and metabolic inhibitor experiment (Fig. 1H-J, S5). The significant increase in the lifetime of free NAD(P)H (*τ*_1_, Fig. S1), suggests a change in the microenvironment (e.g., pH, oxygen) of the free fraction of NAD(P)H that reduces the quenching of the fluorophore. Altogether, the significant changes observed between NAD(P)H and FAD fluorescence lifetime values reflect changes in the microenvironment of the metabolic coenzymes NAD(P)H and FAD and altered metabolic pathway utilizition by quiescent and activated T cells [23–25, 29, 30].

T cells are known to be highly heterogeneous, with phenotypic heterogeneity of surface proteins and effector function observed for CD3^+^CD4^+^ and CD3^+^CD8^+^ T cells [34]. This heterogeneity can arise from 273 the strength of the activating event, the microenvironment of the T cell, and differences in gene regulation at the time of activation [35–37]. Heterogeneity analysis, by heatmaps and histograms, revealed heterogeneous clustering of T cells within the autofluorescence imaging dataset. One of these populations within the quiescent CD3^+^CD8^+^ population, was identified due to a difference in the mean NAD(P)H lifetime which was found to be due to naïve (CD45RO^+^) and memory (CD45RA^+^) CD3^+^CD8^+^ T cells (Fig. 3C-D), which are known to have differing metabolic states: memory T cells have increased glycolytic capacity and mitochondrial mass as compared with naïve T cells [27]. An additional subpopulation was identified within the activated T cell subset and characterized by larger than average cells (Fig. 3B, S6-7). These large cells may be actively dividing cells, a condition which is also accompanied by metabolic and autofluorescence differences [38, 39].

Machine learning approaches are powerful tools for classification of biomedical imaging data and have been used on extracted morphological features of phase-contrast images to identify cancer cells from immune cells, on brightfield images to assess cell cycle, and on phase contrast and autofluorescence images to classify macrophage exposure to LPS [22, 40, 41]. Here, high ROC AUCs (0.95+) were achieved using machine learning techniques to classify T cells as activated or quiescent using the autofluorescence imaging endpoints (optical redox ratio, cell size, NAD(P)H *τ*_*m*_, NAD(P)H *τ*_1_, NAD(P)H *τ*_2_, NAD(P)H *α*_1_, FAD *τ*_*m*_, FAD *τ*_1_, FAD *τ*_2_, and FAD *α*_1_) quantified for each cell. Classification of activation of T cells from CD3^+^CD8^+^ specific isoations was slightly higher than that of T cells from bulk CD3^+^ isolations as might be expected for a homogeneous population (CD3^+^CD8^+^) rather than a heterogeneous population (bulk CD3^+^ populations contain CD4^+^ and CD8^+^ subsets). Although multiple classification models were found to have similar performance, logistic regression was the best fitting model, suggesting that the predicted probability of activation is a linear combination of all 10 of the autofluorescence imaging endpoints. Interestingly, donor normalization (Fig. 2D) of the autofluorescence imaging endpoints did not improve classification accuracy, suggesting that the autofluorescence endpoints reflect changes in T cells with activation that are consistent across donors so generalized models can be used for unspecified donors or patients, which is beneficial for robust implementation of autofluorescence imaging as a universal tool to evaluate T cell activation.

The models for classification of activation in T cells reported here have higher ROC AUC values than the previously reported accuracy of 84-87% found for binary logistic regression classification of morphological and Raman spectra features of control and LPS-exposed macrophages [22]. The increased accuracy obtained in our study could be due to the metabolic information gained from the NAD(P)H and FAD autofluorescence signals, differences in the heterogeneity of the measured populations, and/or differing numbers of cells in the training and testing data sets. Although high classification accuracy was achieved with the machine learning approaches, deep learning methods such as neural networks may achieve improvements in classification accuracy, as has been demonstrated for the classification of cancer cells from immune cells in phase-contrast images [40].

NAD(P)H *α*_1_ was consistently identified as the most important feature for differentiation of quiescent and activated T cells across different feature selection methods (including gain ratio, information gain, *χ*^2^, and random forest), and different subsets of CD3^+^, CD3^+^CD8^+^, and CD69^+^/CD69^−^ T cells (Fig. 2C, S6, S19). The classification analysis also revealed that while models trained on all 10 autofluorescence imaging endpoints yielded the highest accuracy for classification of activation state of T cells, logistic regression using only NAD(P)H *α*_1_ yielded comparably high ROC AUCs and was more accurate for predicting T cell activation than cell size alone (Fig. 2E), or fluorescence intensity measurements (cell size + redox ratio), which can be obtained by wide-field or confocal fluorescence microscopy. Additional label-free methods, including third harmonic generation imaging and Raman spectroscopy of quiescent and activated splenic-derived murine T cells have revealed a significant increase in cell size and lipid content in activated T cells [42]. However, we observed a high variance in cell size within and across patients, which makes it a less important predictor than NAD(P)H lifetime values that change with activation and have lower variance (Fig. S10).

CD3^+^CD4^+^ T cells have a variety of immune-modulating behaviors. While not necessary for activation of CD3^+^CD8^+^ T cells, the presence of CD3^+^CD4^+^ T cells during activation is required for the development 323 of memory CD3^+^CD8^+^ T cells [43]. Additionally, T*REGS* (CD3^+^CD4^+^FoxP3^+^ T cells, 5-10% of peripheral CD3^+^CD4^+^ population) suppress the activation and proliferation of other T cells [44, 45]. Differences in the NAD(P)H and FAD autofluorescence imaging endpoints (Fig. 4, S15) between CD3^+^CD8^+^ T cells cultured with and without CD3^+^CD4^+^ T cells were observed, suggesting autofluorescence imaging is sensitive to CD3^+^CD4^+^ induced changes in CD3^+^CD8^+^ T cells (Fig. 4). However, despite these differences, NAD(P)H *α*_1_ remains the highest weighted feature for classification of activation state (Fig. S16), and activation state of CD3^+^CD8^+^ T cells can be classified from autofluorescence imaging endpoints with high accuracy, regardless of T cell population (Fig. 4D).

Due to the differing physiological functions of CD3^+^CD4^+^ and CD3^+^CD8^+^ T cells [1, 46], it is important to detect CD3^+^CD4^+^ and CD3^+^CD8^+^ subtypes of T cells in addition to the activation state of T cells. Therefore, we explored whether machine learning methods could use autofluorescence imaging data to distinguish between CD3^+^CD8^+^ and CD3^+^CD4^+^ T cells within bulk CD3^+^ populations. Significant 335 differences in NAD(P)H fluorescence lifetime values between CD3^+^CD4^+^ and CD3^+^CD8^+^ T cells suggests variations in metabolic activity upon activation of CD3^+^CD4^+^ and CD3^+^CD8^+^ T cells, which is consistent with previously observed differences in CD3^+^CD4^+^ and CD3^+^CD8^+^ T cell activation: CD3^+^CD4^+^ activation occurs through Myc, ERR*α*, and mTOR, while CD3^+^CD8^+^ T cells activate through Akt and mTOR [47]. These subtle differences in metabolic pathway utilization by CD3^+^CD4^+^ and CD3^+^CD8^+^ T 340 cells enabled high classification accuracy of not only quiescent CD3^+^CD4^+^ from quiescent CD3^+^CD8^+^ cells and activated CD3^+^CD4^+^ from activated CD3^+^CD8^+^ cells, but also all four groups, activated and quiescent CD3^+^CD4^+^ from activated and quiescent CD3^+^CD8^+^ accurately (Fig. 4H). Although successful classification was achieved for CD3^+^CD4^+^ versus CD3^+^CD8^+^ T cells, these changes are much subtler than the metabolic changes with activation, as evidenced by the increased number of cells needed to train the models to achieve high classification accuracy (Fig. 4D,H).

Autofluorescence lifetime imaging has spatial and temporal resolution advantages over traditional assays 347 to survey T cell activation and function. Autofluorescence imaging can be high resolution to allow measurements at the single cell level, allowing insights into metabolic heterogeneity within T cell populations. Additionally, the high spatial resolution and non-destructive nature of autofluorescence imaging maintains 350 the spatial integrity of immune cells, allowing high fidelity measurements on neighboring cells as demonstrated in the combined population of quiescent and activated T cells (Fig. 5A). Finally, autofluorescence imaging also has high temporal resolution (Fig. 5F) allowing time-course study of T cell activation. Alto-353 gether, autofluorescence lifetime imaging of NAD(P)H and FAD of T cells, combined with machine learning for classification, is a powerful tool for non-destructive, label-free assessment of activation status of T cells. NAD(P)H and FAD autofluorescence lifetime imaging is label-free and provides high spatial, temporal, and functional information of cell metabolism, which makes it an attractive tool to evaluate T cells *in vivo* or characterize expanded T cells.

## 4 Methods

### 4.1 T cell Isolation and Culture

This study was approved by the Institutional Review Board of the University of Wisconsin-Madison (#2018-61 0103), and informed consent was obtained from all donors. Peripheral blood was drawn from 6 healthy donors into sterile syringes containing heparin. Two blood draws, 183 days apart, were performed on one donor to evaluate the consistency of the experimental protocol and imaging endpoints. Bulk CD3^+^ T cells or an isolated CD3^+^CD8^+^ T cell subset were extracted from whole blood using negative selection methods (RosetteSep, StemCell Technologies) and cultured in ImmunoCult-XF T cell Expansion Medium (StemCell Technologies). Approximately 24 hours post-isolation, the T cells were divided into two groups, a “quiescent” population that was grown in medium without activating antibodies, and an “activated” population that was cultured in medium supplemented with 25 μl/ml tetrameric antibody against CD2/CD3/CD28 (StemCell Technologies). Quiescent and activated T cell populations were cultured separately for 48 hours at 37°C, 5% CO_2_, and 99% humidity before imaging and subsequent experiments, unless otherwise noted. Prior to imaging, T cells were plated at approximately 200,000 cells/200 μl media on 35 mm poly-d-lysine coated glass bottom dishes (MatTek). To ensure that autofluorescence imaging and the classification models extend for mixed populations of quiescent and activated T cells, a subset of quiescent and activated T cells (48hr of culture with activating antibody) were combined and plated together in a dish 1 hour before imaging.

### 4.2 Autofluorescence Imaging of NAD(P)H and FAD

Fluorescence images were acquired using an Ultima (Bruker Fluorescence Microscopy) two-photon microscope coupled to an inverted microscope body (TiE, Nikon) with an Insight DS+ (Spectra Physics) as the excitation source. A 100X objective (Nikon Plan Apo Lambda, NA 1.3), lending an approximate field of view of 110 μm, was used in all experiments with the laser tuned to 750 nm for NAD(P)H two-photon excitation and 890 nm for FAD two-photon excitation. NAD(P)H and FAD images were acquired sequentially through 440/80 nm and 500/100 nm bandpass filters (Chroma), respectively, by GaAsP photomultiplier tubes (PMTs; H7422, Hamamatsu). The laser power at the sample was 3.0-3.2 mW for NAD(P)H and 4.1-4.3 mW for FAD. Lifetime imaging was performed within Prairie View (Bruker Fluorescence Microscopy) using time-correlated single photon counting electronics (SPC-150, Becker & Hickl, Berlin, Germany). Fluorescence lifetime decays with 256 time bins were acquired across 256×256 pixel images with a pixel dwell time of 4.6 μs and an integration time of 60 s. Photon count rates were ~1-5×10^5^ and monitored during image acquisition to ensure that no photobleaching occurred. The second harmonic generation at 890 nm from red blood cells was used as the instrument response function and had a full width at half maximum of 240 ps. A YG fluorescent bead (*τ* = 2.13 +/− 0.03 ns, n = 6) was imaged daily as a fluorescence lifetime standard [14, 18, 48]. Four to six images per group were acquired.

### 4.3 Antibody Validation

Antibodies against CD4 (clone OKT4, PerCP-conjugated, Biolegend Item #317431, Lot B198303), CD8 (clone SK1, PerCP-conjugated, Biolegend Item #344707, Lot B204988), CD69 (clone FN50, PerCP-conjugated, Biolegend Item #310927, Lot B180058), CD45RA (clone HI100, Alexa 647-conjugated, Biolegend Item #304153, Lot B220325), and CD45RO (clone UCHL1, PerCP-conjugated, Biolegend Item #304251, Lot B219295) were used for validation of cell type and activation. Cells (30,000-200,000 per condition) were stained with 5 μl antibody/106 cells in 50 μl of ImmunoCult-XF T cell Expansion Medium for 30 minutes in the dark at room temperature. Cells were washed with ImmunoCult 1-2 times, resuspended in 50-200 μl of media, and added to the center of a 35 mm poly-d-lysine coated glass bottom dish (MatTek). Cells were kept in a 37°C, 5% CO2, humidified environment until imaging. All cells were imaged within 3 hours of staining. NAD(P)H and FAD fluorescence lifetime images were acquired as described. To identify PerCP positive cells, an additional fluorescence intensity image was acquired with the Titanium:Sapphire laser tuned to 1040 nm and a 690/45 nm bandpass filter before the PMT. For evaluation of Alexa647 fluorescence, the Titanium:Sapphire laser was tuned to 1300 nm for excitation, and a 690/45 nm bandpass filter was used to filter emitted light.

### 4.4 Data Analysis

Fluorescence lifetime decays were analyzed to extract fluorescence lifetime components (SPCImage, Becker & Hickl). A bin of 9 surrounding pixels (3×3) was used to increase the fluorescence counts in each decay. A threshold was used to exclude pixels with low fluorescence signal (i.e. background). Fluorescence lifetime decays were deconvolved from the instrument response function and fit to a 2 component exponential decay model, 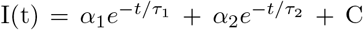, where I(t) is the fluorescence intensity as a function of time, t, after the laser pulse, *α*_1_ and *α*_2_ are the fractional contributions of the short and long lifetime components, respectively (i.e., *α*_1_ + *α*_2_ = 1), *τ*_1_ and *τ*_2_ are the short and long lifetime components, respectively, and 4 C accounts for background light. Both NAD(P)H and FAD can exist in quenched (short lifetime) and unquenched (long lifetime) configurations [9, 13]; therefore, the fluorescence decays of NAD(P)H and FAD are fit to two components.

Images were analyzed at the single cell level to evaluate cellular heterogeneity [49]. NAD(P)H intensity images were segmented into cytoplasm and nucleus using edge detect and thresholding methods in CellProfiler using a customized image processing routine [50]. Images of the optical redox ratio (fluorescence intensity of NAD(P)H divided by the summed intensity of NAD(P)H and FAD) and mean fluorescence lifetimes (*τ*_*m*_ = *α*_1_*τ*_1_ + *α*_2_*τ*_2_) of NAD(P)H and FAD were computed (MATLAB). NAD(P)H and FAD autofluorescence imaging endpoints, including the optical redox ratio, NAD(P)H *τ*_*m*_, NAD(P)H *τ*_1_, NAD(P)H *τ*_2_, NAD(P)H *α*_1_, FAD *τ*_*m*_, FAD *τ*_1_, FAD *τ*_2_, and FAD *α*_1_ were averaged across all pixels within a cell cytoplasm for each segmented cell. Cell size in μm^2^ was also computed from the segmented images using the number of pixels within the 2D-image of the cell * 0.167 μm^2^ (which is the pixel dimension).

Statistical analysis and data representation were performed in R. A generalized linear model was used to evaluate significant differences (*α* = 0.05) of autofluorescence imaging endpoints between quiescent and activated T cells, CD45RA^+^ and CD45RO^+^ cells (Fig. 3), and CD3^+^CD4^+^ and CD3^+^CD8^+^ T cells. Presented boxplots are constructed from the median (central line) and first and third quartiles (lower and upper hinges, respectively). The whiskers extend to the farthest data points that are no further than 1.5* the interquartile range. Dots represent data points beyond 1.5* the interquartile range from the hinge.

### 4.5 Classification

Uniform Manifold Approximate and Projection (UMAP), a dimension reduction technique [26], and z-score heatmaps were used to visualize clustering within autofluorescence imaging data sets (Python and R, respectively). Machine learning classification models and training/testing data sets are summarized in Table S1. Random forest, logistic regression, and support vector machine classification methods were trained to classify activated and quiescent T cells within either the bulk CD3^+^ FLIM data or the isolated CD3^+^CD8^+^ FLIM data (R). For both data sets, gain ratio, *χ*^2^, and random forest feature selection methods were employed to evaluate the contribution of the NAD(P)H and FAD autofluorescence endpoints to the accuracy of classification of quiescent versus activated T cells. These models were trained on data from donors A, B, C, and D because these cells lacked immunofluorescence CD69 validation but were known to be quiescent or activated by culture conditions (n = 4131 CD3^+^ cells, n = 2655 CD3^+^CD8^+^ cells). Models were tested on data from T cells from donors B, E, and F with CD69 immunofluorescence validation of activation state (n = 696 CD3^+^ cells, n = 595 CD3^+^CD8^+^ cells). Random forest models were developed to classify CD3^+^CD4^+^ from CD3^+^CD8^+^ T cells, and cells were randomly assigned to training and test data sets for a range of train/test proportions from 12.5% to 87.5%. Each model was replicated 50 times with new training and test data generated before each iteration. Logistic regression models were also estimated for the classification of T cell activation from imaging endpoints of combined quiescent and activated CD3^+^ T cells (both conditions together within the images). Observations were randomly divided into training and testing data sets (90%/10%, respectively), and presented ROC curves are the average of 1000 iterations of randomly selected training and testing data.

### 4.6 Seahorse Assay

Quiescent and activated T cells were plated at 5×10^6^ cells/ml on a Seahorse 96-well plate in unbuffered RPMI medium without serum. Oxygen consumption rate (OCR) and extracellular acidification rate (ECAR) measurements were obtained every 6.5 minutes for 5 cycles. A generalized linear model was used to determine statistical significance (*α* = 0.05) within OCR and ECAR measurements between control and activated T cells.

### 4.7 Metabolic Inhibitors

Quiescent and activated (48 hr) CD3^+^ T cells were plated on poly-d-lysine coated 35 mm glass bottom dishes at a concentration of ~200,000 cells/200 μl ImmunoCult T cell Expansion Medium as previously described (T cell Isolation and Culture). The metabolic inhibitors antimycin A (1 μM), rotenone (1 μM), 2-deoxy-d-glucose (2DG, 50 mM), Bis-2-(5-phenylacetamido-1,3,4-thiadiazol-2-yl)ethyl sulfide (BPTES, 20 μM), and 5-(Tetradecyloxy)-2-furoic acid (TOFA, 50 1 μg/ml) were added singly, except for antimycin A and rotenone which were added together, to the dishes prior to imaging. Cells were incubated with antimycin A and rotenone for ten minutes, 2DG for ten minutes, BPTES for 1 hour, and TOFA for 1 hour. Fluorescence lifetime images of NAD(P)H and FAD were acquired for 6 random fields of view as described above. A generalized linear model was used to determine autofluorescence imaging endpoints with statistical significance (*α* = 0.05) between control and inhibitor-exposed cells.

### 4.8 Activation Time Course

Quiescent CD3^+^ T cells were isolated and plated for imaging as previously described. NAD(P)H lifetime images were acquired as described but with an image size of 128×128 pixels and an integration time of 15 s. Images were acquired sequentially for 2 minutes (8 frames), then 5 μl PBS was added to the cells as 473 a mock treatment, and NAD(P)H fluorescence lifetime images were acquired for 10 minutes (40 frames). Subsequently, 5 μl of activating tetrameric antibody (anti-CD2/CD3/CD28) was added and NAD(P)H fluorescence lifetime images were acquired for 10 minutes (40 frames). NAD(P)H FLIM images were analyzed in SPCImage as described. Individual cells and cell compartments (nucleus, cytoplasm) were manually segmented (author I.J.), and the autofluorescence imaging endpoints were averaged across all pixels within the segmented region (ImageJ). This procedure was repeated for 3 dishes for a total of 34 analyzed cells.

### 4.9 Data Availability

The datasets generated during and/or analyzed during the current study are available from the corresponding authors on reasonable request.

### 4.10 Code Availability

All code and algorithms generated during the current study are available from the corresponding authors on reasonable request.

## 5 Acknowledgments

The authors would like to thank Arezoo Movaghar for insightful discussions of feature selection and machine learning classification methods and Rebecca Schmitz for her assistance with formatting of paper figures. This work was funded by the NIH NCI R01 CA205101 (to M.C.S); the Biotechnology Training Program of the National Institute of General Medical Sciences of the National Institutes of Health, award #T32GM008349 (to K.S.); NIH awards R01DK098672 and P41GM108538 (to D.J.P.) and T32DK007665 (to N.M.N.); the NSF Graduate Research Fellowship Program, DGE-1747503 (to K.M); and the National Science Foundation under Grant No. EEC-1648035 (to K.S.).

## 6 Author Contributions

AW and MS conceived the central hypotheses, and KM contributed the hypothesis on distinguishing CD3^+^CD8^+^ naïve versus memory T cell autofluorescence properties. KM and AW designed and performed the experiments with assistance from NP. AW and IJ analyzed the data. NN and KM performed the Seahorse assay. CW provided statistical insight and data analysis code. KS and MS supervised the project. AW wrote the initial draft of the manuscript. All authors contributed to data interpretation and the final manuscript.

## 8 Correspondence

Correspondence to Alex J. Walsh or Melissa C. Skala.

