## Supplemental Tables and Figures for "Label-free Method for Classification of T cell Activation"

Table S1: Machine learning classification models

| Classification | Models | Train/Test Data* | N | Iterations | Figure |
|---|---|---|---|---|---|
| Bulk CD3$^+$ Isolation: Quiescent vs. Activated | Random forest<br>SVM<br>Logistic Regression | 85.6% status known by culture conditions/14.4% with CD69 validation | 4827 cells<br>6 donors | NA | 2D-E |
| CD3$^+$CD8$^+$ Isolation: Quiescent vs. Activated | Random forest<br>SVM<br>Logistic Regression | 81.7% status known by culture conditions/18.3% with CD69 validation | 3250 cells<br>6 donors | NA | 2D, 2F |
| Bulk CD3$^+$ Isolation, CD3$^+$CD8$^+$ Subset: Quiescent vs. Activated | Random forest | 12.5%/87.5%<br>25%/75%<br>37.5%/62.5%<br>50%/50%<br>62.5%/37.5%<br>75%/25%<br>87.5%/12.5% | 253 cells<br>3 donors | 50 | 4D |
| CD3$^+$CD8$^+$ Isolation, CD3$^+$CD8$^+$ Subset: Quiescent vs. Activated | Random forest | 12.5%/87.5%<br>25%/75%<br>37.5%/62.5%<br>50%/50%<br>62.5%/37.5%<br>75%/25%<br>87.5%/12.5% | 213 cells<br>3 donors | 50 | 4D |
| Bulk CD3$^+$ Isolation: Quiescent CD4$^+$ vs. Quiescent CD8$^+$ | Random forest | 12.5%/87.5%<br>25%/75%<br>37.5%/62.5%<br>50%/50%<br>62.5%/37.5%<br>75%/25%<br>87.5%/12.5% | 149 cells<br>3 donors | 50 | 4H |
| Bulk CD3$^+$ Isolation: Activated CD4$^+$ vs. Activated CD8$^+$ | Random forest | 12.5%/87.5%<br>25%/75%<br>37.5%/62.5%<br>50%/50%<br>62.5%/37.5%<br>75%/25%<br>87.5%/12.5% | 434 cells<br>3 donors | 50 | 4H |
| Bulk CD3$^+$ Isolation: Activated CD4$^+$ vs. Activated CD8$^+$ vs. Quiescent CD4$^+$ vs. Quiescent CD8$^+$ | Random forest | 12.5%/87.5%<br>25%/75%<br>37.5%/62.5%<br>50%/50%<br>62.5%/37.5%<br>75%/25%<br>87.5%/12.5% | 583 cells<br>3 donors | 50 | 4H |
| Bulk CD3$^+$ Isolation, Combined CD69$^+$/CD69$^-$ T cells: CD69$^-$ vs. CD69$^+$ | Logistic regression | 90%/10% | 250 cells<br>1 donor | 1000 | 5E |

* Cells randomly assigned unless otherwise noted.

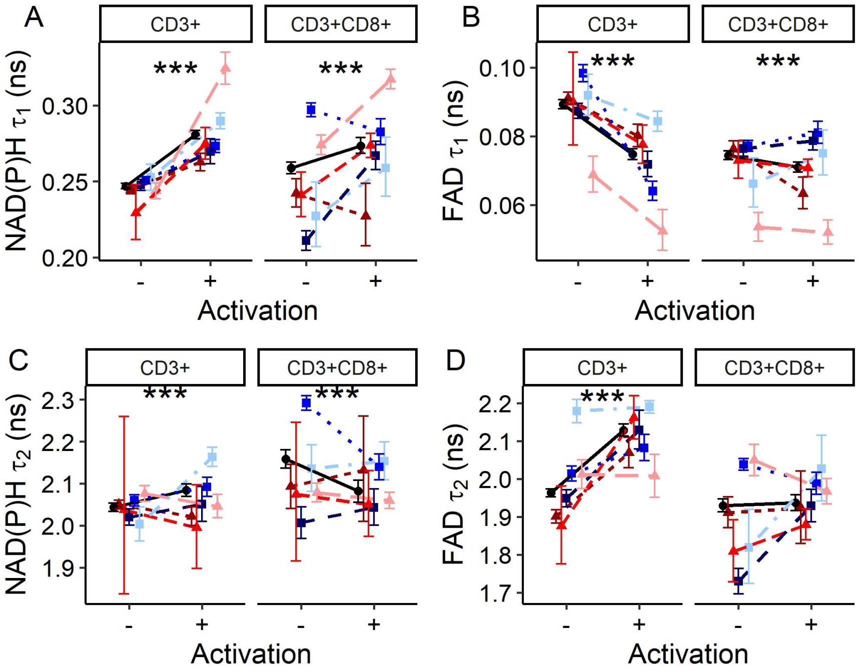


Figure S1: **NAD(P)H and FAD fluorescence lifetime components reveal metabolic differences with activation in bulk CD3$^+$ and isolated CD3$^+$CD8$^+$ T cells.** (A) NAD(P)H $\tau_1$, (B) FAD $\tau_1$, (C) NAD(P)H $\tau_2$, and (D) FAD $\tau_2$ of quiescent and activated CD3$^+$ and CD8$^+$ T cells. Black circles represent mean of all data (6 donors), triangles (donors A [dark red], B [medium red], and F [light red]) represent data from female donors, squares (donors C [dark blue], D [medium blue], and E [light blue]) represent data from male donors. Each color shade represents data from an individual donor. Data are mean +/- 99% CI. *** $p<0.0001$.

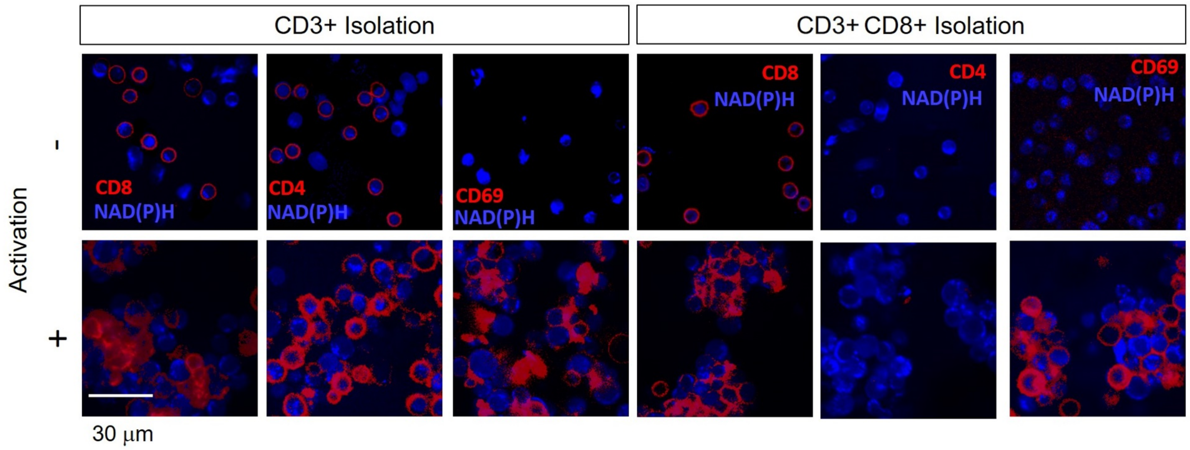


Figure S2: **Immunofluorescence staining of CD4, CD8, and CD69 verifies T cell subtype and activation.** Fluorescence images of quiescent and activated CD3$^+$ (right) and CD3$^+$CD8$^+$ (left) T cells stained with antibodies for CD4, CD8, or CD69 overlaid on NAD(P)H fluorescence intensity images.

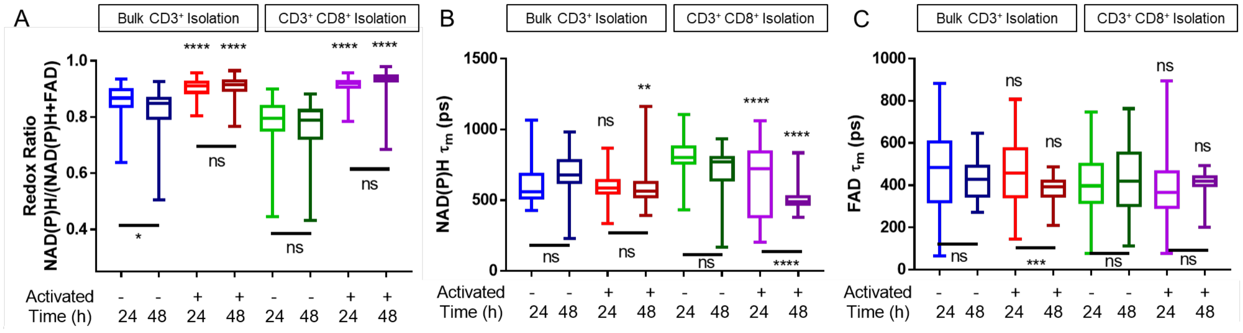


Figure S3: **NAD(P)H and FAD autofluorescence endpoints of bulk CD3$^+$ and isolated CD3$^+$CD8$^+$ T cells are consistent for cells activated for 24 or 48 hours.** Stars above indicate signficance between quiescent and activated populations within a time point (24 or 48 hr). Bars with stars indicate p-values between groups imaged at 24 and 48 hr of activation. ns= not signficant, * $p<0.05$, ** $p<0.01$, *** $p<0.001$, **** $p<0.0001$. 1 donor n=44-209 cells/group from one donor.

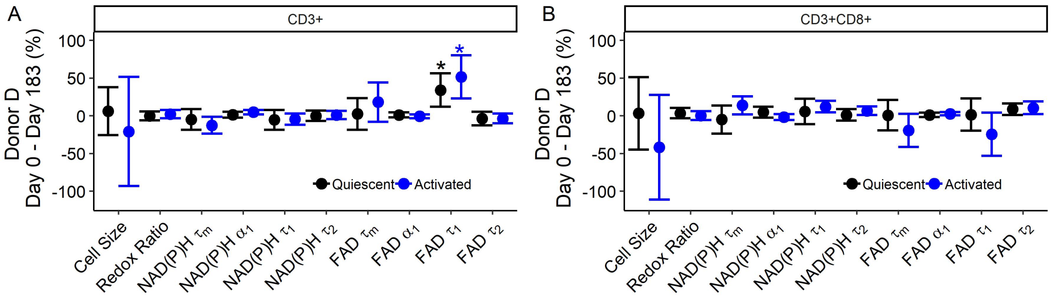


Figure S4: **Autofluorescence imaging endpoints are consistent across two different isolations from blood drawn from the same donor.** Percent difference between mean autofluorescence imaging endpoints of CD3$^+$ (A) and CD3$^+$CD8$^+$ (B) T cells from two blood draws (183 days apart) from the same donor. Data are mean +/- 95% CI. * $p<0.05$.

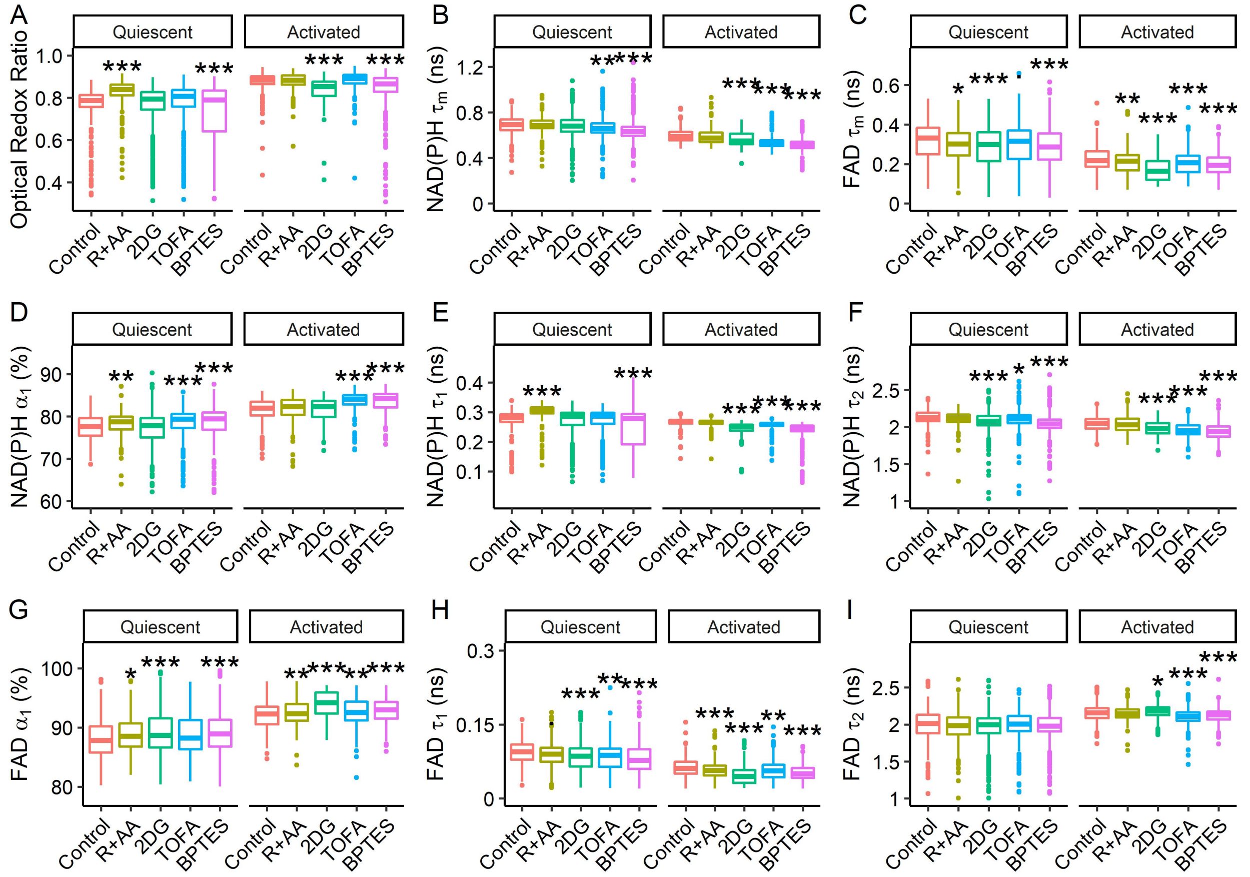


Figure S5: **Autofluorescence imaging of NAD(P)H and FAD resolves metabolic perturbations in quiescent and activated CD3$^+$ T cells**. Quiescent and activated bulk CD3$^+$ T cells were incubated with the metabolic inhibitors rotenone and antimycin A (R+AA, electron transport chain inhibitors), 2-de-oxy-D-glucose (2DG, glycolysis inhibitor), 5-(Tetradecyloxy)-2-furoic acid (TOFA, fatty acid synthesis inhibitor), and Bis-2-(5-phenylacetamido-1,3,4-thiadiazol-2-yl)ethyl sulfide (BPTES, glutaminolysis inhibitor). n = 165-558 cells, * $p<0.05$, ** $p<0.01$,*** $p<0.001$, Generalized linear model.

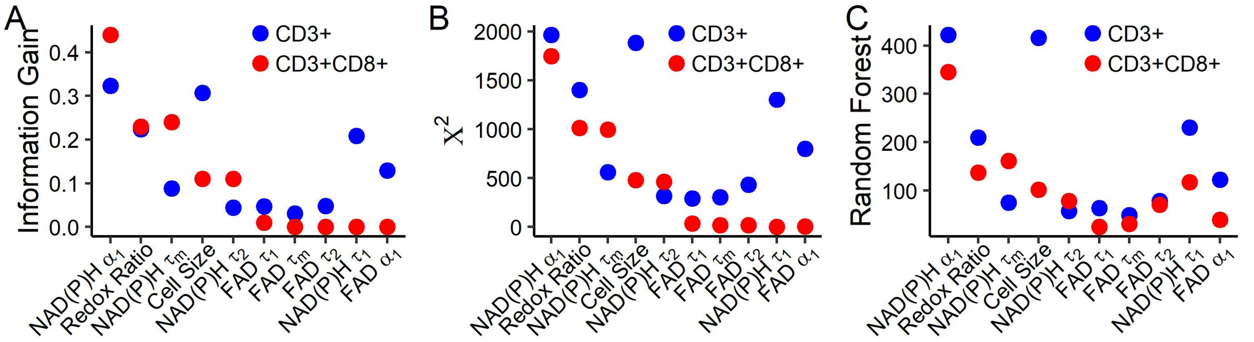


Figure S6: **Relative importance of autofluorescence endpoints for classification of activation status.** Feature weights determined for bulk CD3$^+$ and isolated CD3$^+$CD8$^+$ T cells by (A) information gain, (B) $\chi^2$, and (C) random forest trees.

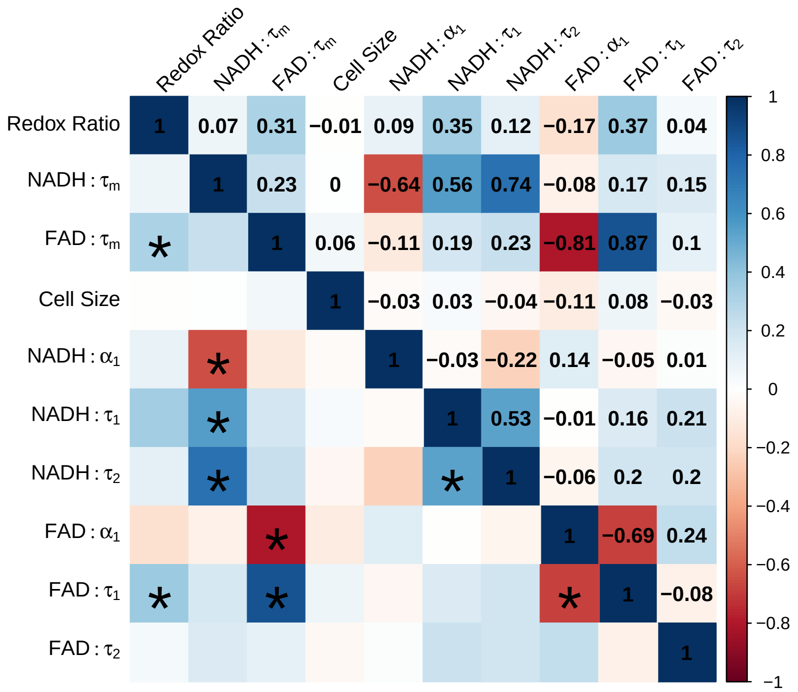

Figure S7: **Correlation matrix for NAD(P)H and FAD fluorescence lifetime imaging endpoints.** Boxes are color-coded to Spearman's correlation coefficient, averaged for quiescent $CD3^+$ T cells, activated $CD3^+$ T cells, quiescent $CD3^+CD8^+$ T cells, and activated $CD3^+CD8^+$ T cells. Values are Spearman's correlation coefficient. * signifies p-values less than 0.001.

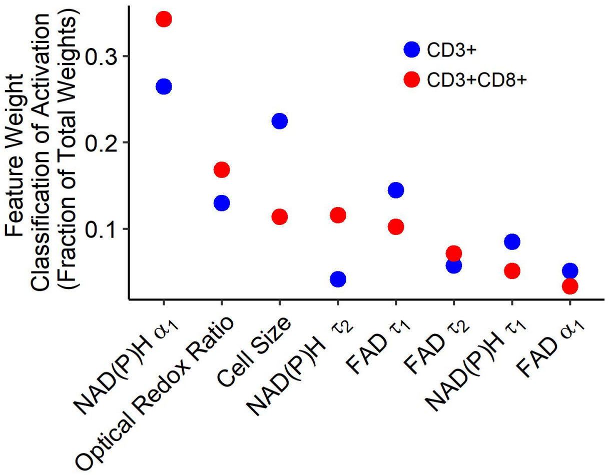


Figure S8: **NAD(P)H $\alpha_1$ is the most weighted feature for classification of activation of T cells.**Normalized feature weight of NAD(P)H and FAD autofluorescence imaging endpoints, excluding the composite endpoints, NAD(P)H $\tau_m$ and FAD $\tau_m$, as determined from random forest trees for the classification of activation of bulk $CD3^+$ and isolated $CD3^+CD8^+$ T cells.

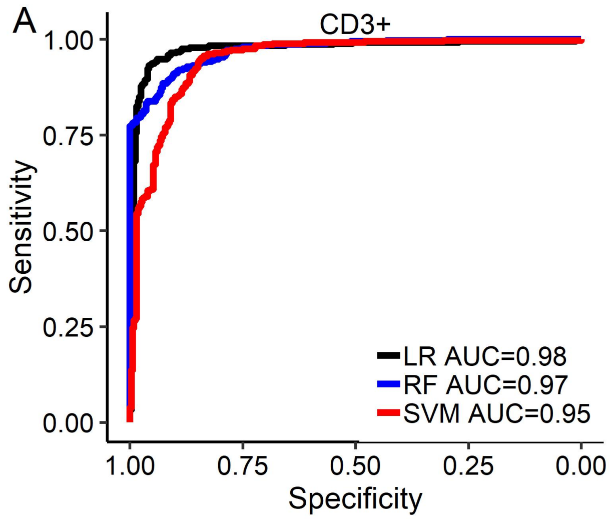


Figure S9: **Similar performance of multiple models is achieved for classification of T cell activation.** Logistic regression, random forest, and support vector machine models using all 10 autofluorescence imaging endpoints for classification of activation status in (A) bulk $CD3^+$ and (B) isolated $CD3^+CD8^+$ T cells.

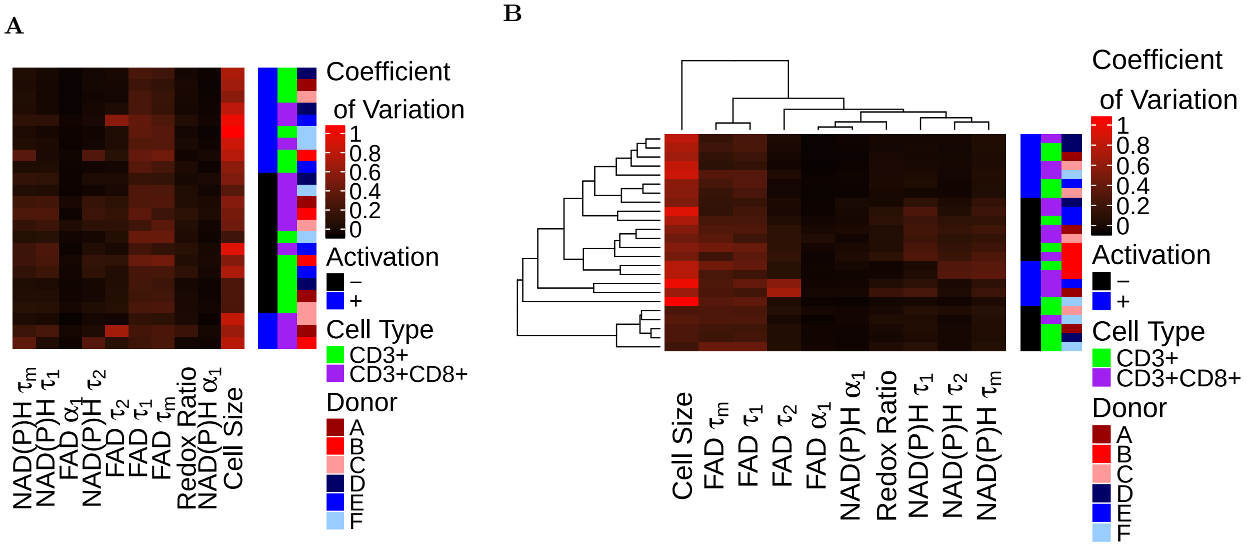


Figure S10: **Coefficient of variation heatmaps reveal variance within endpoints across T cell groups.** Coefficient of variation heatmap with each row representing the data from one donor, cell type, and activation state. Rows and columns are clustered by (A) the z-score dendrogram from Fig. 3A or (B) by a dendrogram created from clustering of the coefficient of variation.

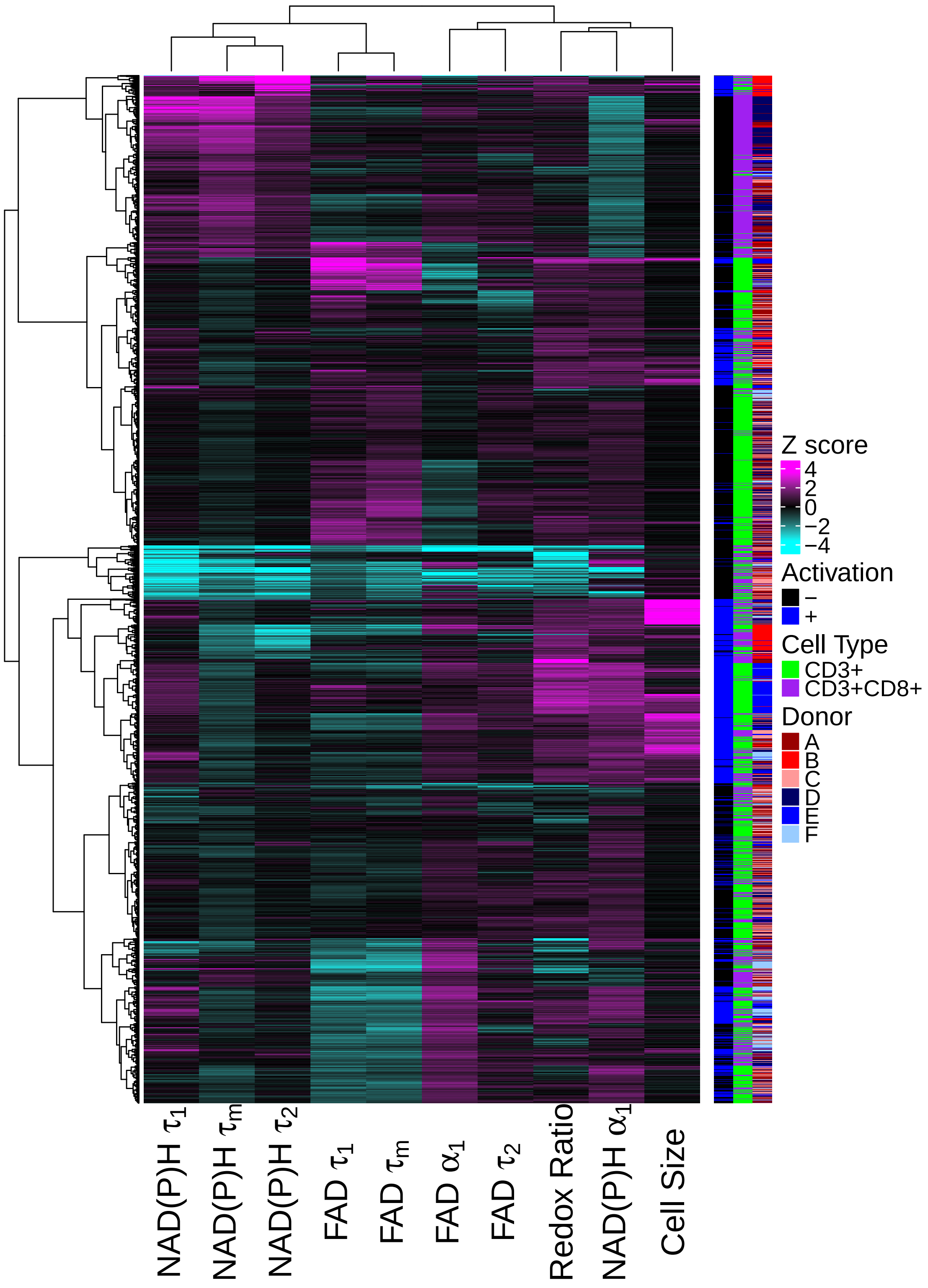


Figure S11: **Autofluorescence imaging reveals T cell heterogeneity.** Heatmap of z-scores of autofluorescence endpoints of quiescent and activated T cells. Each row is one cell, data from all cells from 6 donors. T cells cluster by activation status and cell isolation (bulk $CD3^+$ or isolated $CD3^+CD8^+$) but not by donor.

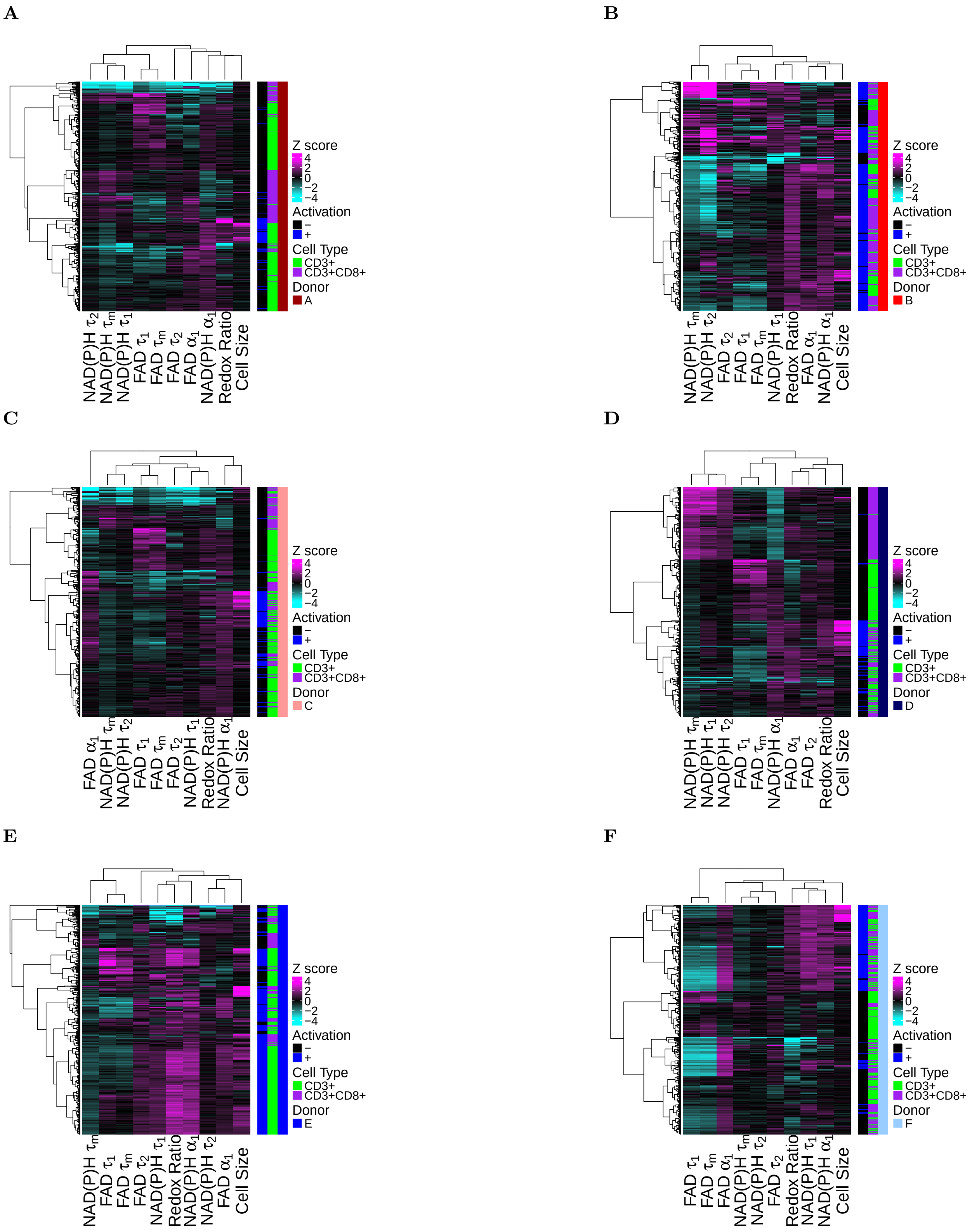


Figure S12: **Autofluorescence imaging reveals heterogeneity within donors.** Heatmap of z-scores of autofluorescence endpoints of all quiescent and activated bulk $CD3^+$ and isolated $CD3^+CD8^+$ T cells (rows) measured for each of the 6 donors.

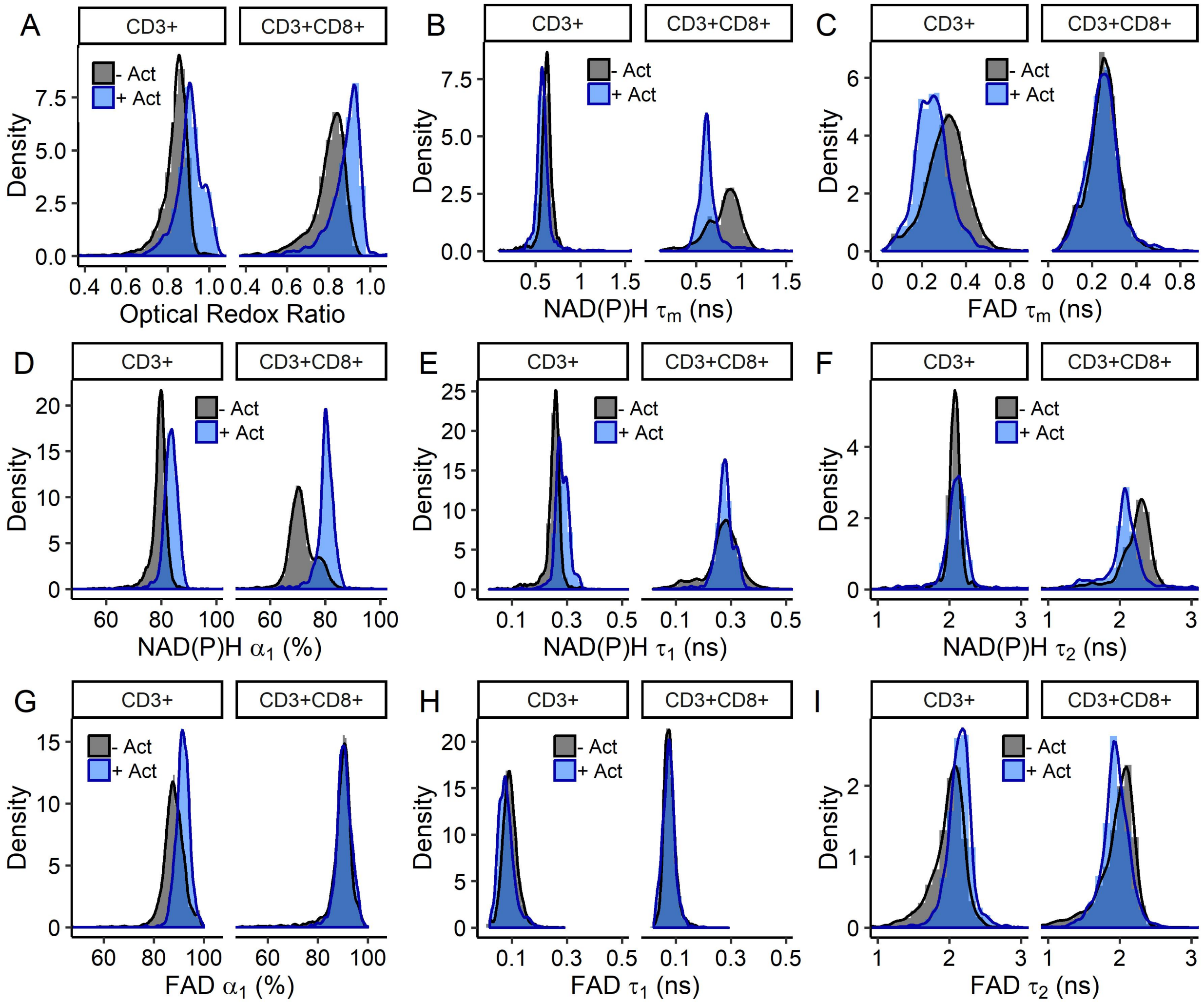


Figure S13: **Histograms of autofluorescence endpoint frequencies of quiescent (black) and activated (blue) bulk $CD3^+$ and isolated $CD3^+CD8^+$ T cells.** Data is pooled from all donors and corresponds to Fig. 1B-G and S1.

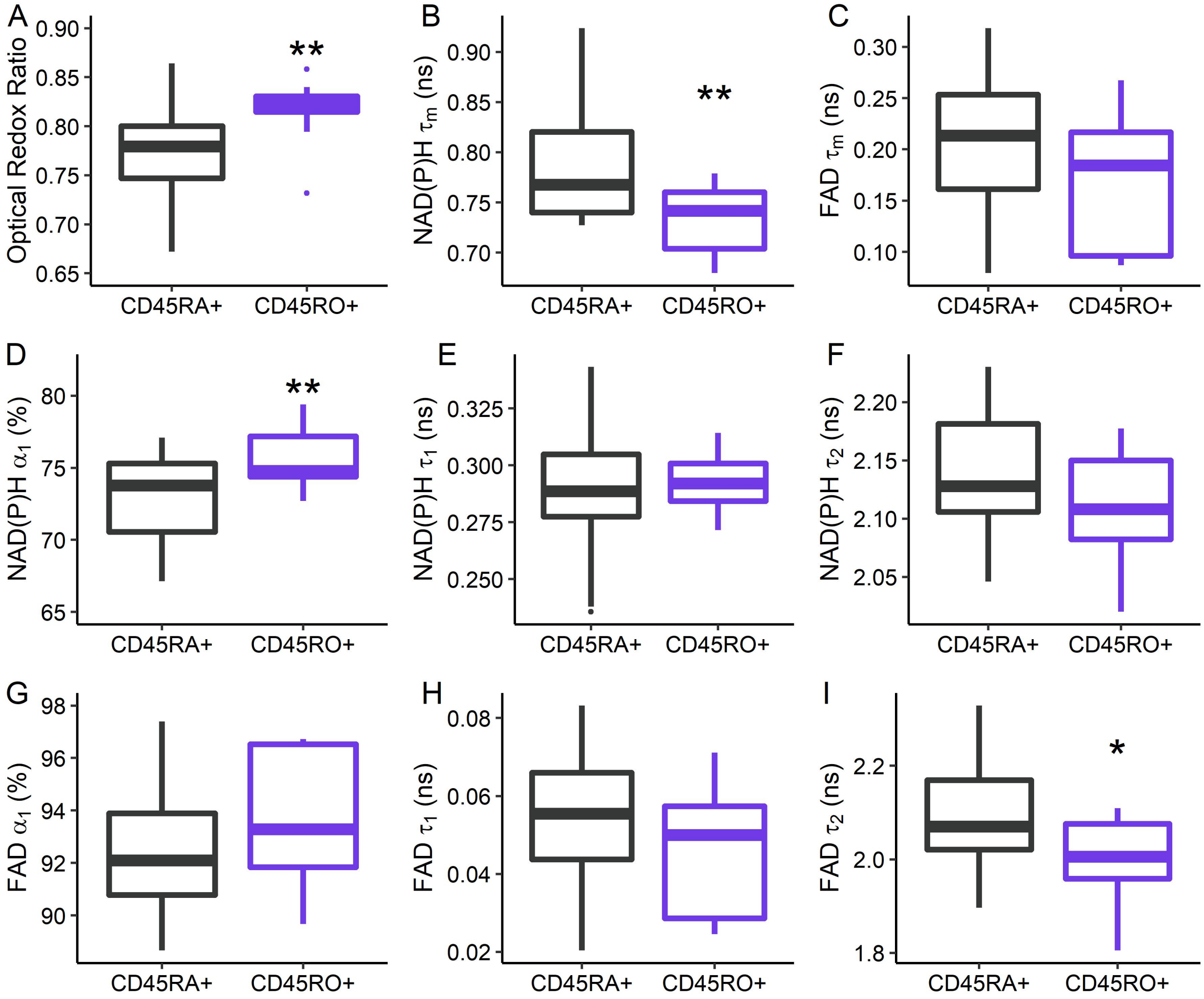


Figure S14: **NAD(P)H and FAD autofluorescence endpoints resolve metabolic differences between memory (CD45RO$^+$, n=11 cells) and naïve (CD45RA$^+$, n=27 cells) CD3$^+$CD8$^+$ T cells.** * $p<0.05$, ** $p<0.01$, generalized linear model

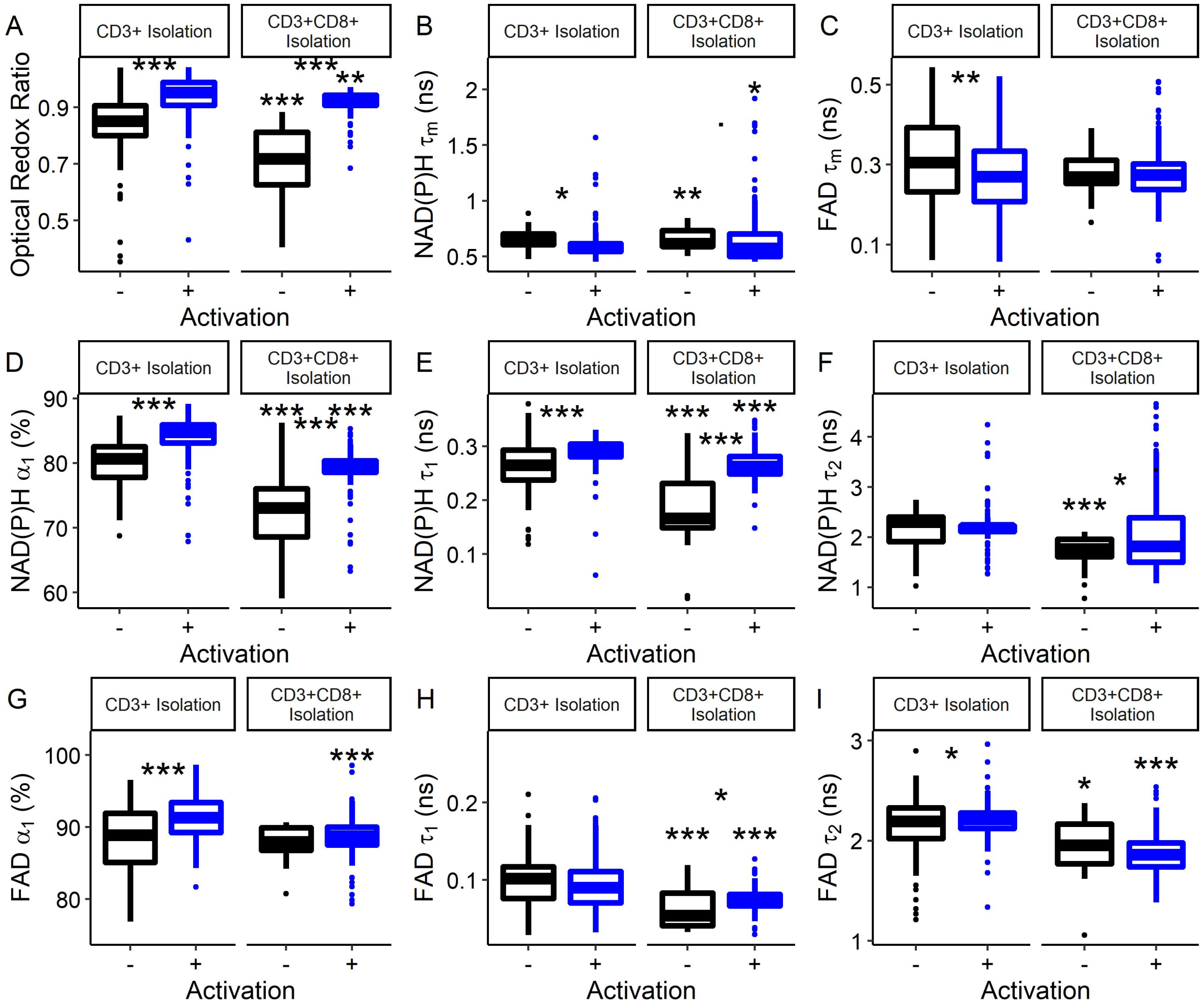


Figure S15: **NAD(P)H and FAD autofluorescence endpoints reveal metabolic differences between CD3$^+$CD8$^+$ cells from the bulk CD3$^+$ isolation and CD3$^+$CD8$^+$ cells from the CD3$^+$CD8$^+$-specific isolation.** * $p<0.05$, ** $p<0.01$, *** $p<0.001$, generalized linear model. Stars between quiescent and activated boxplots compare quiescent and activated CD3$^+$CD8$^+$ T cells within an isolation (CD3$^+$ or CD3$^+$CD8$^+$). Stars above the quiescent box plot represent signficance between quiescent CD3$^+$CD8$^+$ T cells from the bulk CD3$^+$ and CD3$^+$CD8$^+$-specific isolations. Stars above the activated box plot represent signficance between activated CD3$^+$CD8$^+$ T cells from the bulk CD3$^+$ and CD3$^+$CD8$^+$-specific isolations. n=20-434 cells from 1 donor.

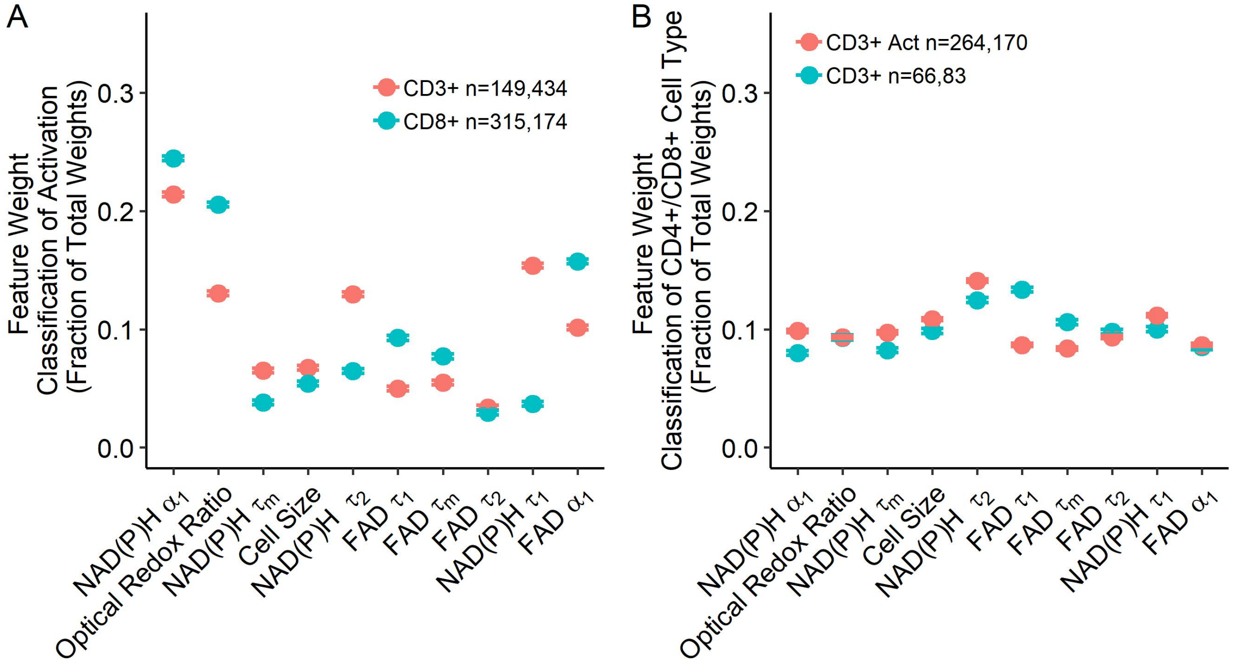


Figure S16: **Autofluorescence feature importance for classification of activation status and $CD3^+CD4^+/CD3^+CD8^+$ T cells.** (A) Normalized feature weights as determined from random forest trees for the classification of activation of T cells from bulk $CD3^+$ (n=149 $CD69^-$ cells and n=434 $CD69^+$ cells) and $CD3^+CD8^+$-specific (n=315 $CD69^-$ cells and n=174 $CD69^+$ cells) isolations. (B) Normalized feature weights as determined from random forest trees for classification of quiescent $CD3^+CD4^+$ (n=264) from quiescent $CD3^+CD8^+$ (n=170) T cells within bulk $CD3^+$ isolations, and normalized feature weights for classification of activated $CD3^+CD4^+$ (n=66) from activated $CD3^+CD8^+$ (n=83) T cells within bulk $CD3^+$ isolations.

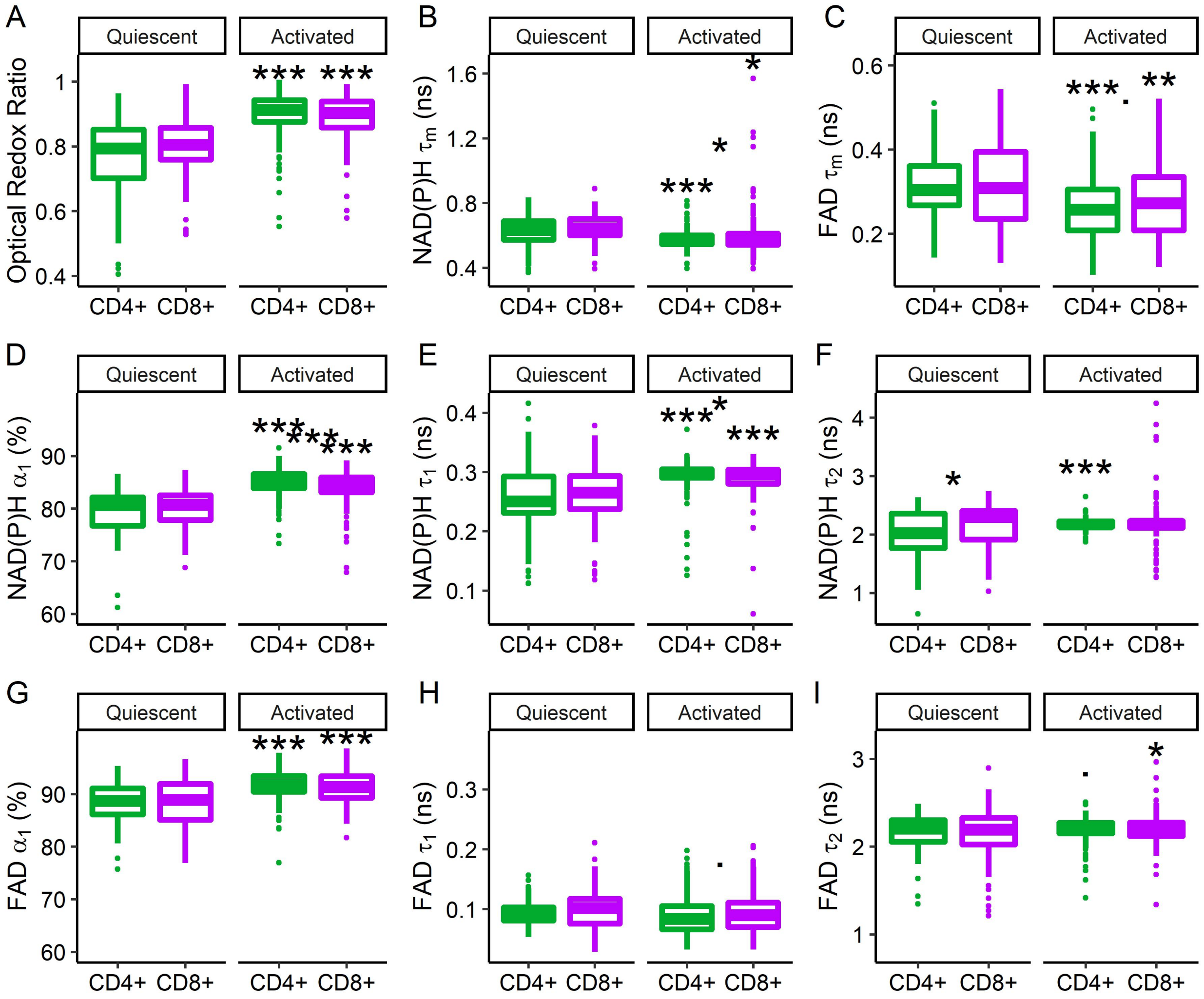


Figure S17: **NAD(P)H and FAD autofluorescence imaging endpoints of $CD3^+CD4^+$ and $CD3^+CD8^+$ T cells within bulk $CD3^+$ isolations.** * $p<0.05$, ** $p<0.01$, *** $p<0.001$ generalized linear model. Stars between $CD3^+CD4^+$ and $CD3^+CD8^+$ boxplots represent significance between $CD3^+CD4^+$ and $CD3^+CD8^+$ T cells. Stars above the activated $CD3^+CD4^+$ or $CD3^+CD8^+$ boxplot indicate signficance between quiescent and activated $CD3^+CD4^+$ or $CD3^+CD8^+$ T cells, respectively. n=94-541 cells. $CD4^+$ and $CD8^+$ expression was verified by same-cell immunofluorescence.

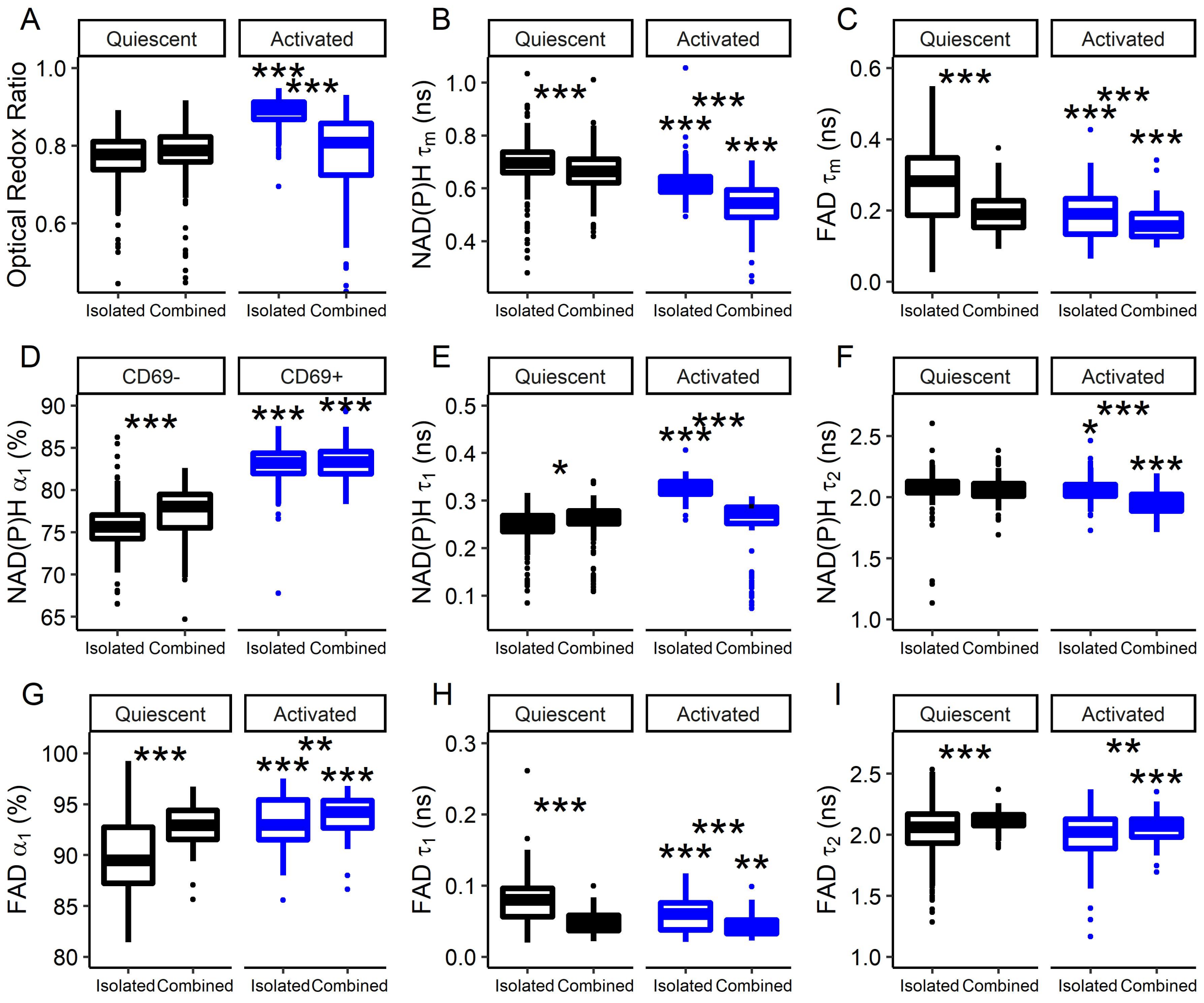


Figure S18: **Mixed culture of quiescent and activated T cells affects T cell autofluorescence.** NAD(P)H and FAD autofluorescence imaging endpoints of mixed $CD3^+$ T cells within isolated cell populations and mixed activation ($CD69^+/CD69^-$) populations. Subsets of $CD3^+$ T cells were cultured in the absence (quiescent) or presence (activated) of a tetrameric antibody against CD2/CD3/CD28 for 48 hours. Quiescent and activated cells were first imaged separately ("isolated") and then combined into one population ("together"), which was imaged 1 hour after combining cells. ** $p<0.01$, *** $p<0.001$ generalized linear model. Stars between Isolated and Together boxplots represent significance between $CD3^+$ T cells cultured as isolated $CD69^-/CD69^+$ populations and the corresponding $CD69^-/CD69^+$ cells of the "Together" population. Stars above the $CD69^+$ Isolated or Together boxplot indicate signficance between $CD69^-$ and $CD69^+$ $CD3^+$ T cells from Isolated or Together populations, respectively. n=289-438 cells from one donor. CD69 expression was used as a marker of activation and was verified by same-cell immunofluorescence.

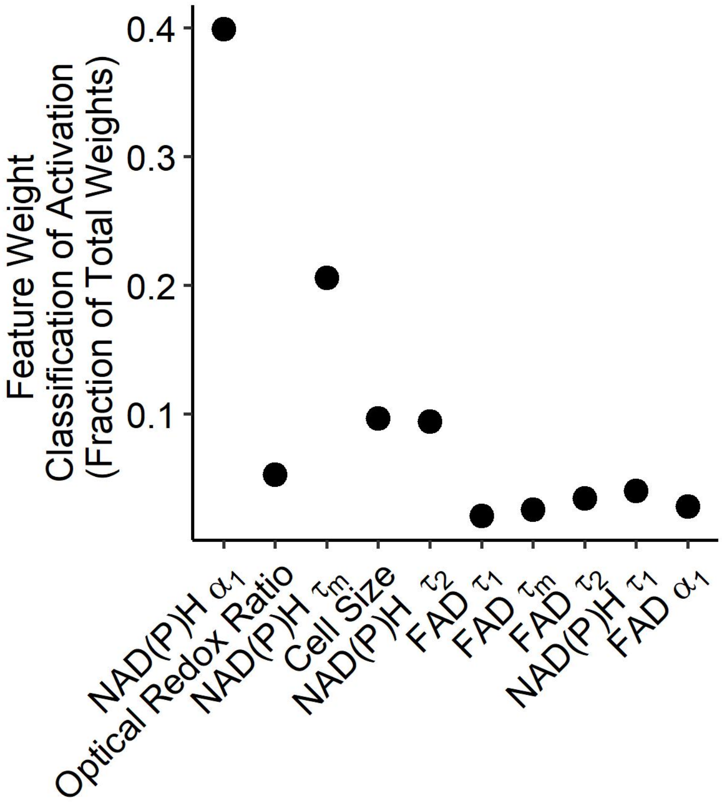


Figure S19: **NAD(P)H $\alpha_1$ is the highested weighted feature for the classification of activated or quiescent T cells within a combined population of T cells.**Normalized feature weight of NAD(P)H and FAD autofluorescence imaging endpoints as determined from random forest trees for the classification of activation of combined quiescent and activated $CD3^+$ T cells.
